# An *in vivo* Dissection, and Analysis of Socio-Affective Symptoms related to Cerebellum-Midbrain Reward Circuitry in Humans

**DOI:** 10.1101/2023.09.29.560239

**Authors:** Linda J. Hoffman, Julia M. Foley, Josiah K. Leong, Holly Sullivan-Toole, Blake L. Elliott, Ingrid R. Olson

**Affiliations:** Temple University, Department of Psychology and Neuroscience, Philadelphia, PA, USA; University of Arkansas, Department of Psychological Science, Fayetteville, AR, USA

**Author notes:** Address correspondence to: Linda J. Hoffman, Department of Psychology, Temple University, 1701 N. 13^th^Street, Philadelphia, PA 19122.

**Keywords:** cerebellum, dopamine, ventral tegmental area, diffusion imaging, depression, psychosis, reward

## Abstract

Emerging research in non-human animals implicates cerebellar projections to the ventral tegmental area (VTA) in appetitive behaviors, but these circuits have not been characterized in humans. Here, we mapped cerebello-VTA white-matter connectivity in humans using probabilistic tractography on diffusion imaging data from the Human Connectome Project. We uncovered the topographical organization of these connections by separately tracking from parcels of cerebellar lobule VI, crus I/II, vermis, paravermis, and cerebrocerebellum. Results revealed that connections from the cerebellum to the VTA predominantly originate in the right hemisphere, interposed nucleus, and paravermal cortex, and terminate mostly ipsilaterally. Paravermal crus I sends the most connections to the VTA compared to other lobules. We discovered a medial-to-lateral gradient of connectivity, such that the medial cerebellum has the highest connectivity with the VTA. Individual differences in microstructure were associated with measures of negative affect and social functioning. By splitting the tracts into quarters, we found that the socio-affective effects were driven by the third quarter of the tract, corresponding to the point at which the fibers leave the deep nuclei. Taken together, we produced detailed maps of cerebello-VTA structural connectivity for the first time in humans and established their relevance for trait differences in socio-affective regulation.

## 1. Introduction

The cerebellum is often overlooked by research on affective and motivated behaviors. However, over the last twenty years, neuroimaging studies have found cerebellar alterations in a range of neuropsychiatric disorders such as depression (Phillips, Hewedi, Eissa, & Moustafa, 2015), schizophrenia (Andreasen & Pierson, 2008; Moberget et al., 2018), autism spectrum disorder (ASD) (Courchesne, Yeung-Courchesne, Press, Hesselink, & Jernigan, 1988; Wang, Kloth, & Badura, 2014), and substance abuse (Miquel, Gil-Miravet, & Guarque-Chabrera, 2020; Moulton, Elman, Becerra, Goldstein, & Borsook, 2014). For instance, a recent study examined the new incidence of psychiatric disorders following the onset of cerebellar disease in older adults (Delle Chiaie et al., 2015). Results show that bipolar disorder was diagnosed in 31% of patients with cerebellar disease, highlighting the potential importance of the cerebellum in emotion regulation. Converging evidence from volumetric studies has found changes in cerebellar thickness, density, or volume in individuals with bipolar disorder relative to healthy controls (Baldacara et al., 2011; DelBello, Strakowski, Zimmerman, Hawkins, & Sax, 1999; Eker et al., 2014; Kim et al., 2013; Lippmann et al., 1982; Lupo et al., 2021; Mills, Delbello, Adler, & Strakowski, 2005; Nasrallah, McCalley-Whitters, & Jacoby, 1982) (see also Brambilla, Barale, Caverzasi, and Soares (2002); Laidi et al. (2015); Monkul et al. (2008)).

Mood disorders can be conceptualized as a problem in reward seeking, or motivated behavior(Gonen, Sharon, Pearlson, & Hendler, 2014; Russo & Nestler, 2013). New research has shown that the cerebellum may play an essential role in motivated behavior through its influence on the ventral tegmental area (VTA), a midbrain nucleus that plays a key role in motivated behavior as the origin of many dopaminergic cell bodies(Novello, Bosman, & De Zeeuw, 2022). Monosynaptic connectivity between the cerebellum and VTA has been established from axon tracer studies (Judd, Lewis, & Person, 2021; Perciavalle, Berretta, & Raffaele, 1989; Phillipson, 1979; R. S. Snider, Maiti, & Snider, 1976; Watabe-Uchida, Zhu, Ogawa, Vamanrao, & Uchida, 2012) in cats and rodents as well as optogenetic investigations in mice (Baek, Park, Kim, Yamamoto, & Tanaka-Yamamoto, 2022; Carta, Chen, Schott, Dorizan, & Khodakhah, 2019). In a seminal study, Carta et al. (2019) transiently silenced the cerebellum-VTA pathway in mice and found that stimuli that were usually rewarding—other mice—became uninteresting after the pathway was optogenetically inactivated. A following study by Low et al. (2021) found that upregulating the pathway in mice could increase feeding behavior, while downregulating the pathway decreased feeding, regardless of baseline satiety. Moreover, selectively activating, or silencing neurons projecting from the cerebellum could alter VTA activity such that more or less dopamine was released (Low et al., 2021). These findings suggest that portions of the cerebellum regulate reward signals that catalyze or inhibit reward-driven behavior, even without direct manipulation of central reward regions.

These findings suggest that the cerebello-VTA pathway may play an essential, yet overlooked, role in motivated behavior. Moreover, this circuit may be relevant to understanding the pathophysiology of diverse socio-affective disorders with disrupted reward-seeking behavior including disorders with prominent reductions in social motivation (e.g., social anxiety, schizophrenia, autism spectrum disorder (ASD) (Berridge & Robinson, 2016), eating and substance use disorders (Berridge & Robinson, 2016; Morales & Berridge, 2020), and mood disorders (Pizzagalli, 2014; Watson & Naragon-Gainey, 2010). To date, no research has characterized the cerebello-VTA structural connectivity in humans.

Here we asked the foundational question: does the cerebello-VTA pathway exist in humans? We performed diffusion tractography from the deep cerebellar nuclei and cerebellar cortex to visualize this pathway for the first time in humans. We then mapped the laterality and topographical organization of the projections from the cerebellum. We restricted our analysis to cerebellar lobules VI, crus I and II, which are the only regions in the posterior lobe that have vermal regions corresponding to the neocerebellum (Amore et al., 2021), and which converge with the cerebellar regions linked functionally to reward processing (Popal, 2023). Finally, we tested the prediction that individual difference in cerebello-VTA tract microstructure would be associated with variability in social and affective function.

## 2. Results

Due to the large number of analyses in the current report, a narrative summary of the statistical results is provided below. To view detailed information on raw streamline analyses, including Bonferroni-corrected p-values, bootstrapped confidence intervals, effect sizes, and effect size magnitude summaries, please refer to **Tables 1** and **2**. For the success rate of all tractography reconstructions, please see **Table S3**. Finally, companion tables of length- and waypoint-corrected results can be found in supplemental information (**figure S1**, as well as **tables S4-S5** and **S6-S7**, respectively). In the following sections we present findings from the cerebellar cortex to the VTA using the DCN as waypoints (=polysynaptic tracts) and from the DCN to the VTA (=monosynaptic tracts).

**Table 1.**
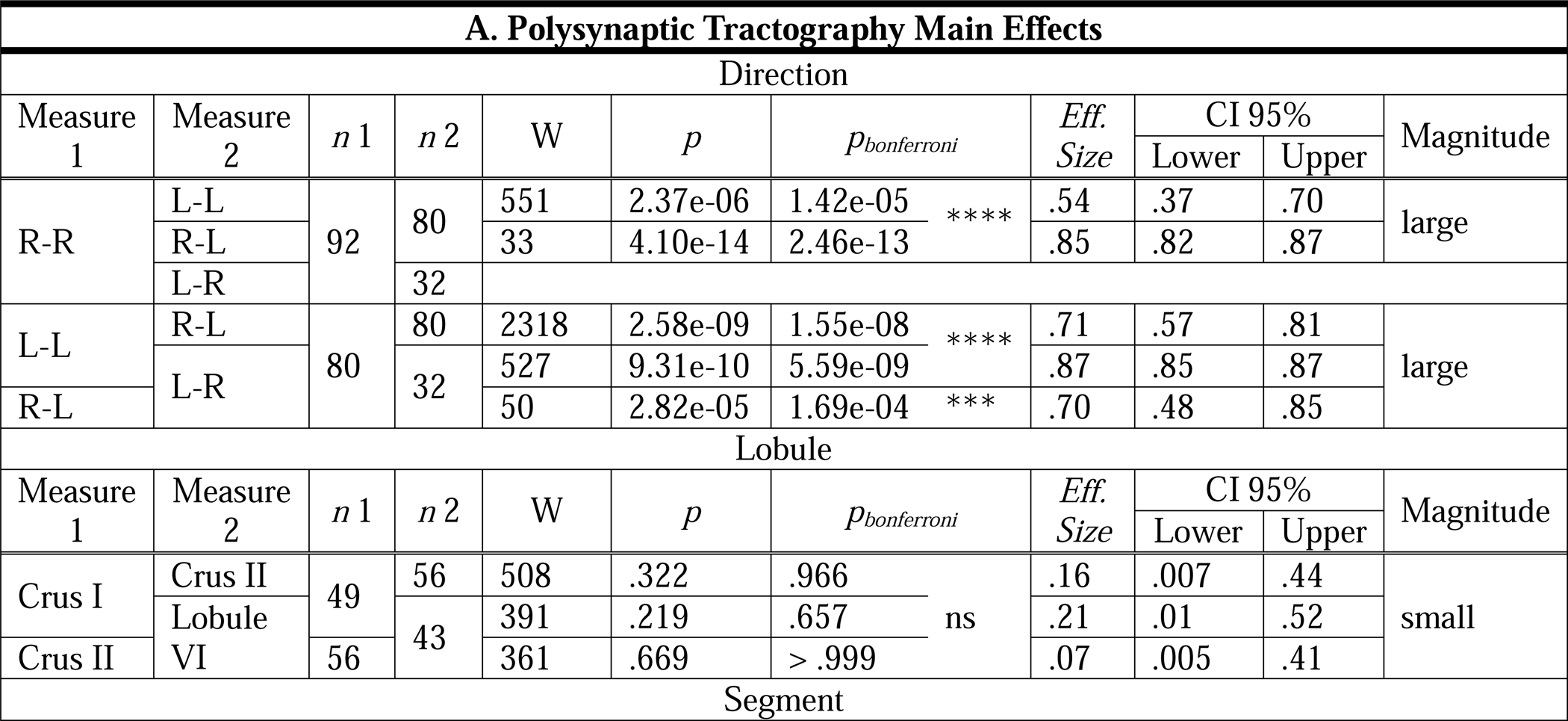
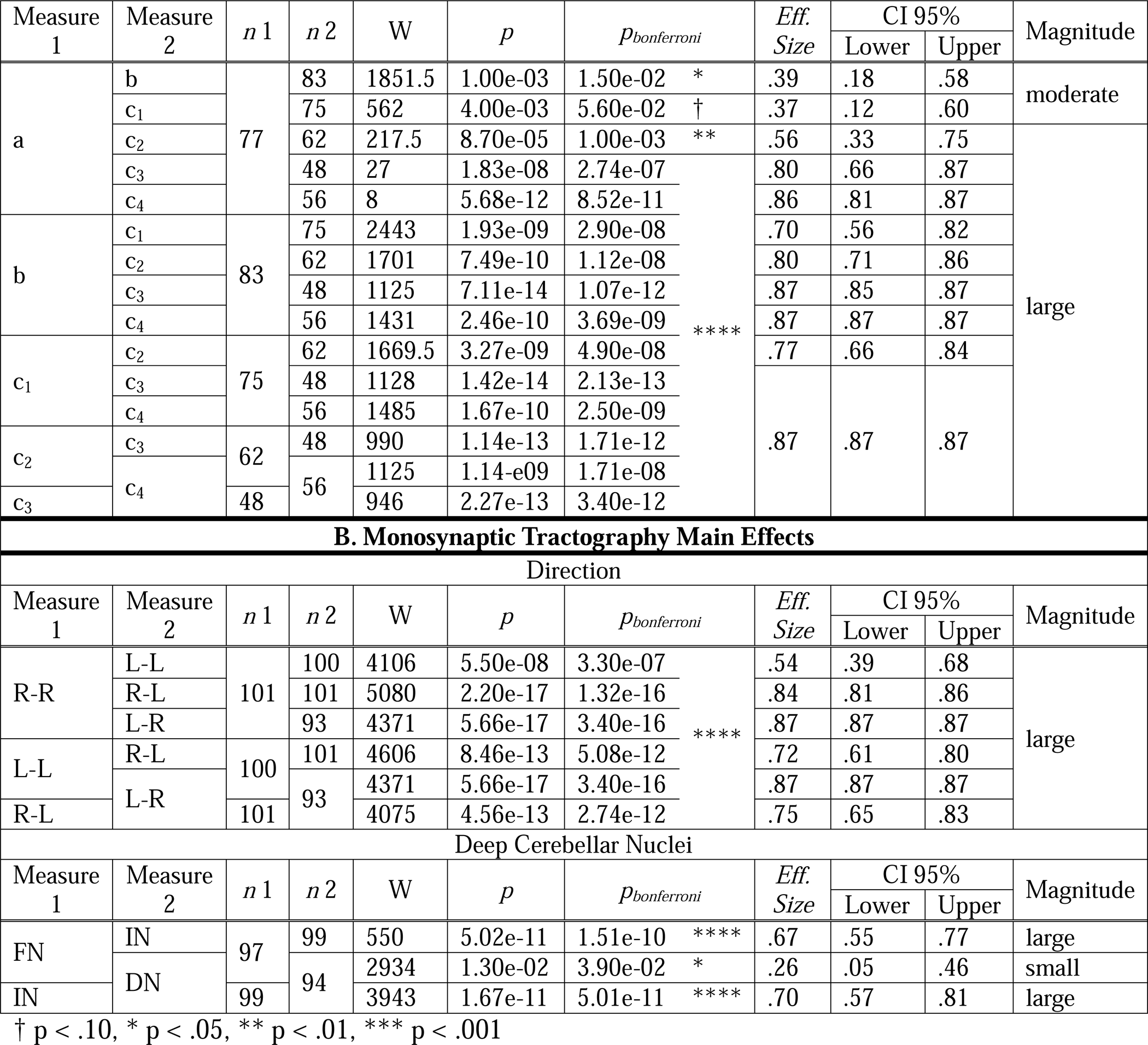
**(A)** Nonparametric pairwise comparisons using Wilcoxon sign-rank test investigating streamline count differences based on tract direction, lobule, and segment. All confidence intervals were bootstrapped with 10,000 iterations. **(B)** Repetition of these analyses to investigate streamline count differences based on tract direction and deep nucleus for monosynaptic tractographies. For all analyses, confidence intervals were bootstrapped with 10,000 iterations. In cases where all instances of a tract were larger than its comparison tract, inferential statistics are not reported. Directions are coded as follows: R-R = right to right (ipsilateral tract), L-L = left to left (ipsilateral tract), R-L = right to left (contralateral tract), L-R = left to right (contralateral tract). Segments are coded as follows: a = vermis, b = paravermis; c_1_ – c_4_ = neocerebellum, with higher numbers indicating more lateral segments. FN=fastigial nucleus; IN=interposed nucleus; DN = dentate nucleus.

**Table 2.**
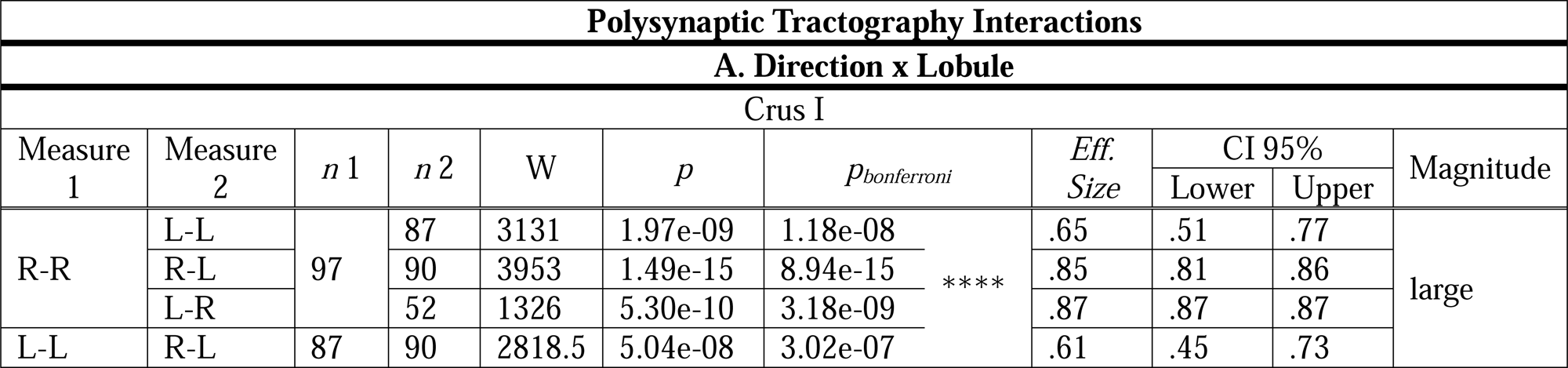
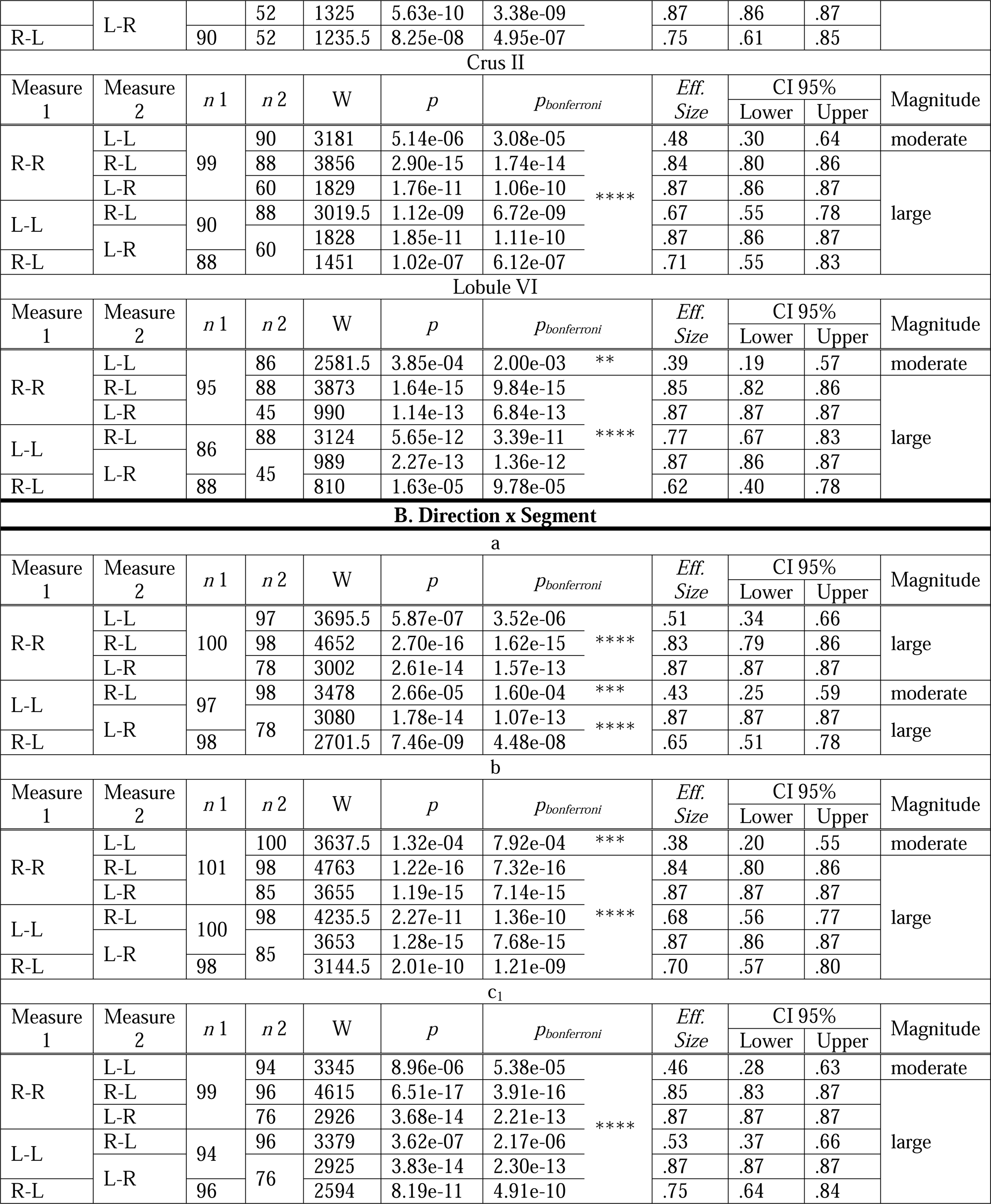
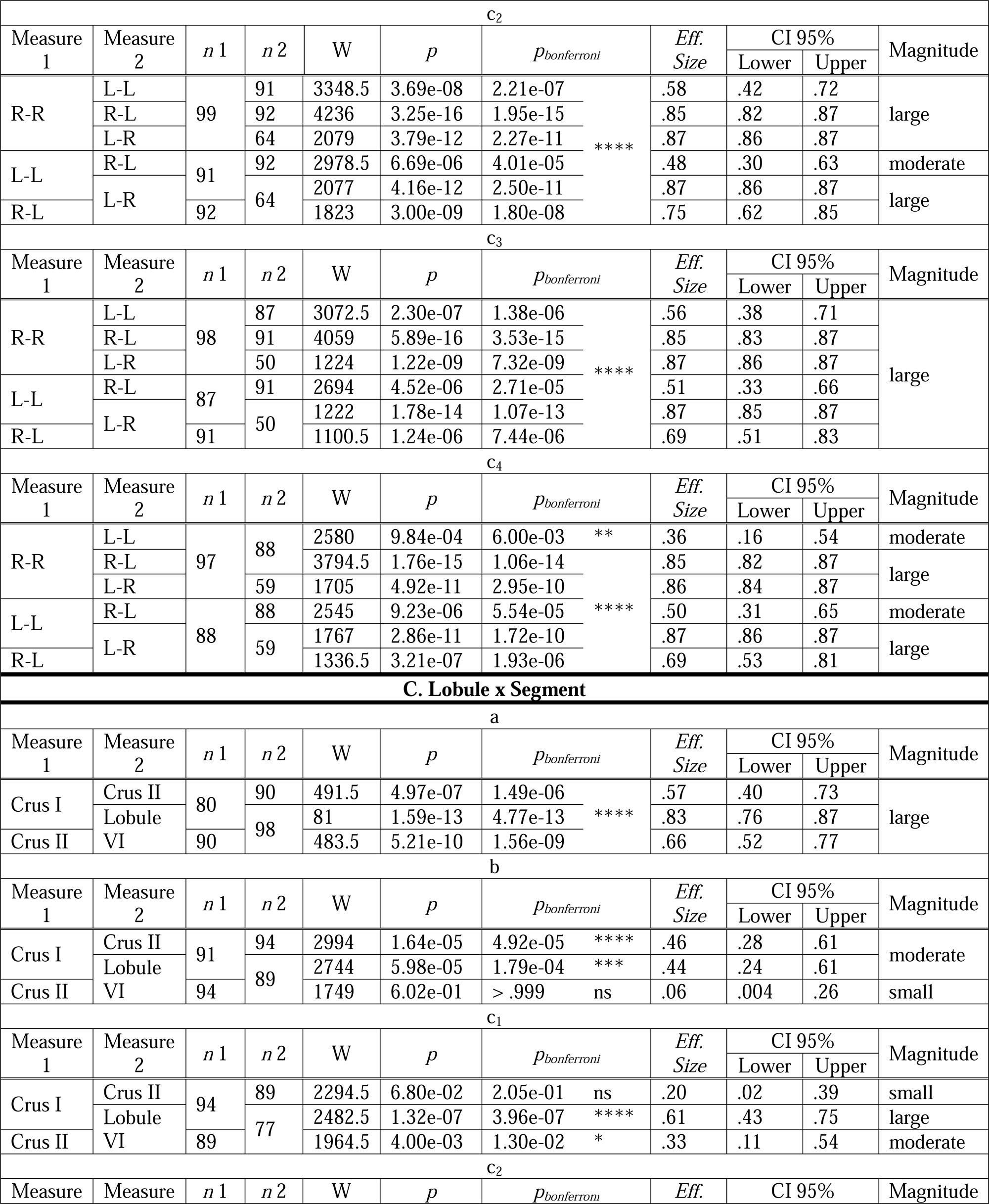
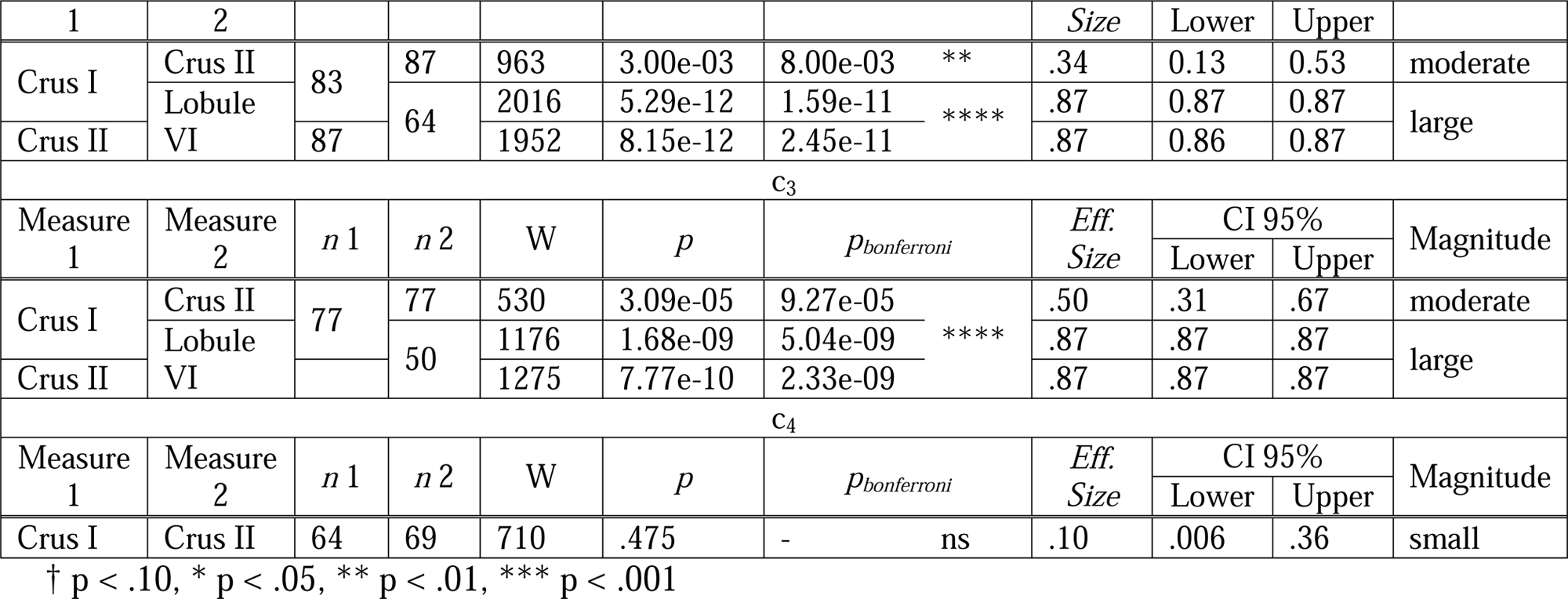
Nonparametric pairwise comparisons using Wilcoxon signed-rank test investigating interactions between **(A)** direction and lobule; **(B)** direction and segment; and **(C)** lobule and segment. All confidence intervals are bootstrapped using 10,000 iterations to yield stable estimates around the effect size.

### 2.1. Anatomy findings

#### 2.1.1. Main Effects

##### Polysynaptic tracts

First, focused Wilcoxon signed-rank tests revealed a main effect of direction such that more streamlines travel from the cerebellar cortex to the ipsilateral VTA. All possible direction combinations were significantly different from one another. There was an effect of laterality such that there was stronger connectivity from the right cerebellum to the VTA. This holds for both the ipsilaterally and contralateral (e.g., decussating) tract. This effect persists after controlling for DCN volume and tract length.

Second, pairwise comparisons revealed no main effect of lobule. This finding held when controlling for average tract length. However, when controlling for DCN volume as a confound, we found a significant difference in connectivity between lobule VI and crus I, as well as a trending difference (after Bonferroni correction) between lobule VI and crus II. Both effects are of moderate magnitude and suggest higher streamline density coming from lobule VI as compared to crus I/II.

Third, there is a main effect of “segment” such that nearly all segments significantly differ from one another. Pairwise comparisons between every segment revealed a higher number of streamlines coming from the paravermis (segment b) compared to the vermis proper. From there, there is a medial to lateral gradient of decreasing connectivity. However, when controlling for average tract length as a potential confound, the difference between the vermis (a) and paravermis (b) becomes non-significant, and the difference between the vermis and c_1_ becomes significant such that the vermis has more connectivity to the VTA overall, while all other comparisons remain effectively unchanged. Finally, after controlling for DCN volume, the difference in direction between the vermis (a) and paravermis (b) flips, such that the vermally-seeded tracts contain higher streamline density travelling through the fastigial nucleus than does the paravermally-seeded tracts travelling through the interposed nucleus. Moreover, when controlling for DCN size, the difference between the vermis and c_1_ remains significant. All other findings remain unchanged when controlling for DCN waypoint volume, meaning that the gradient of lessening connectivity with more laterally positioned segments endures.

##### Monosynaptic tracts

Focused Wilcoxon signed-rank tests revealed a main effect of direction such that more streamlines travel from the DCN to the ipsilateral VTA. All possible direction combinations were significantly different from one another. There was an effect of laterality such that there was stronger connectivity from the right DCN to the VTA. This holds for both the ipsilaterally and contralateral (e.g., decussating) tract. This effect persists after controlling for tract length.

Second, there is a main effect of DCN. Pairwise comparisons revealed that most streamlines going to the VTA come from the interposed nucleus, second most from the fastigial, and the least coming from the dentate. This effect was not confounded by tract length.

All findings concerning main effects are summarized in **Figure 1**.

**Figure 1.**
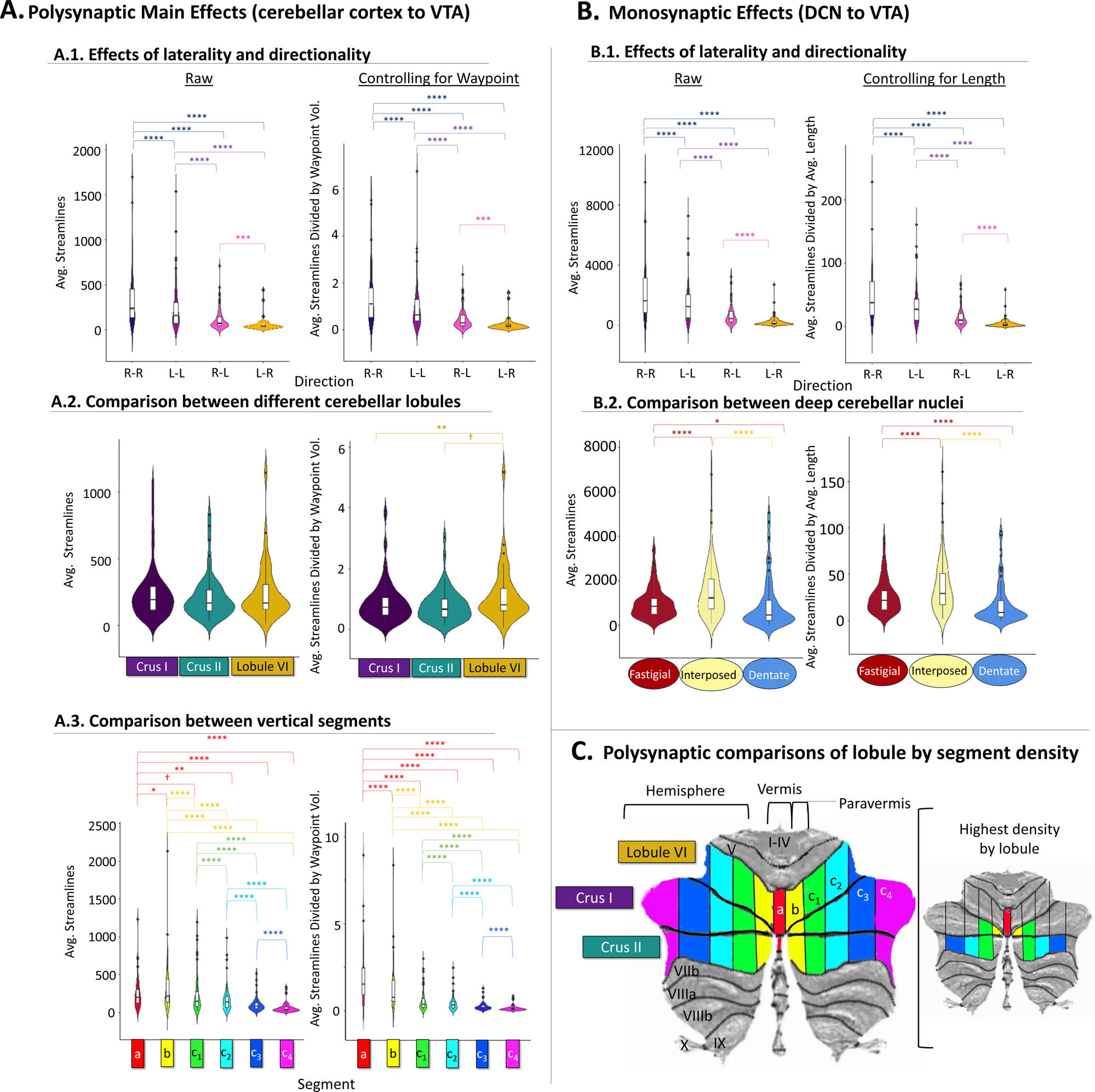
A summary of statistical results investigating gross pairwise comparisons. These are reported for both the polysynaptic (A) and monosynaptic (B) reconstructions. Each result is illustrated twice: on the left are the results using raw streamlines, and on the right are the streamline results when controlling for waypoint volume in voxels. The results for the lobule by segment interactions are illustrated by cerebellar flat maps in section (C). The flat map on the left is meant to orient the viewer to the location of the cerebellar lobules and segments under study, while the flat map on the right has highlighted on the lobule for each segment where streamline density was the highest. Note that segment c_1_ is highlighted in crus I and II as both lobules contained more streamlines in this segment than lobule VI without being statistically different form each other. Also, segment c_4_ contains no remaining highlights since the streamline count in crus I and II did not significantly differ from each other in this segment (not this segment did not extend into lobule VI). **Significance Flags**: † p < .10, * p < .05, ** p < .01, *** p < .001

#### 2.1.2. Interactions

For the polysynaptic tractographies, there was neither an interaction between lobule and direction, nor between segment and direction. That is to say that regardless of the lobule or segment that was seeded, the ipsilateral tract always contained more streamlines to the VTA than the contralateral tract, and the tract seeded in the right cerebellum always contained more VTA connectivity than those seeded in the left. These findings are robust and persist when controlling for both length and waypoint volume in the case of the polysynaptic reconstructions.

There were, however, interactions between segment and lobule, such that different segments showed differing levels of connectivity between the three lobules under study. This is important because it provides insights into which lobule provides the most connectivity to the VTA within each segment. These findings are detailed below and in **Figure 2**. Findings for length and waypoint-controlled lobule by segment comparisons are summarized in **Supplemental Figure 1B**.

**Figure 2.**
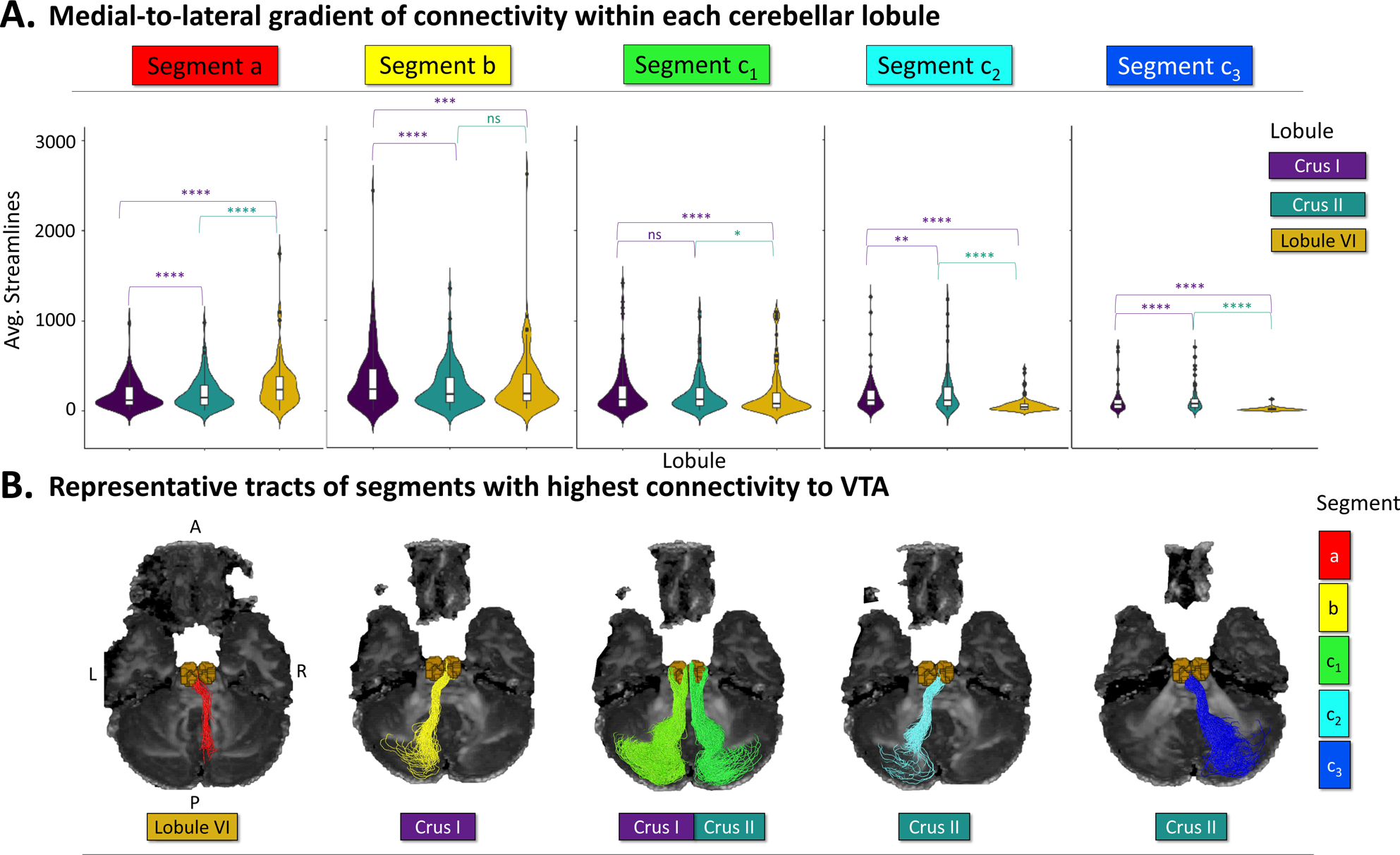
Topographical organization of cerebellum-VTA connectivity. A. Medial segments of each cerebellar lobule have the highest connectivity with the VTA. B. Representative tractography reconstructions for the segment of each cerebellar lobule with the highest connectivity to the VTA. Note that the laterality and directionality of the tracts are not meaningful but show instances of decussation the polysynaptic reconstructions. **Significance Flags**: † p < .10, * p < .05, ** p < .01, *** p < .001

##### Vermis (Segment a)

All possible lobule comparisons were significantly different from each other in the vermis, such that crus I vermal connectivity to the VTA contained the lowest connectivity, while lobule VI contained the most. This interaction holds after controlling for both average tract length and waypoint volume. This means that within the vermal regions under study, lobule VI contains the highest contribution of connections to the VTA, a finding that holds after holding multiple confounds constant.

##### Paravermis (Segment b)

There was a significant difference between paravermal connectivity to the VTA originating in crus I such that most of the connectivity from this segment originated in this lobule as compared to crus II and lobule VI. Furthermore, there was no difference between the relative amount of paravermal connectivity coming from crus II and lobule VI. This finding held when controlling for volume of the DCN waypoints. However, when controlling for average streamline length, only crus I and crus II remained significantly different from one another, with this difference remaining in the same direction.

##### Cerebrocerebellum Segment c_1_

There was an interaction between lobule and segment in the case of c_1_, such that lobule VI contains fewer VTA projections than both crus I and crus II while the latter two lobules remain non-significantly different from each other. This effect persists when controlling for waypoint volume. However, when controlling for length, it emerges that crus I contains more VTA connectivity than crus II and lobule VI, while the difference between crus II and lobule VI becomes non-significant.

##### Cerebrocerebellum Segments c_2_ & c_3_

All possible lobule comparisons in segment c_2_ and c_3_ were significantly different from one another, such that crus II contains the most VTA connectivity in these segments, while lobule VI contains the least. These effects held after controlling for average tract length and waypoint volume within these segments.

##### Cerebrocerebellum Segment c_4_

There was no difference in segment c_4_ between crus I/II, and one did not emerge when controlling for average tract length or waypoint volume.

### 2.2. Brain-Behavior Findings

Using the segments of highest density identified in the anatomical investigation, we examined the relationship between tract microstructure and behavior. Robust correlations were revealed between all and self-report measures; they are summarized in **Table 3**, and supplemental **Table S8**, respectively. Descriptive statistics and FA profiles are found in supplemental **Table S9**, **S10, and S12**. A complete correlation matrix with all brain, self-report, and control variables are reported in supplemental **Table S11**. For all brain-self-report analyses, we further split the tracts into quarters, corresponding to the more superficial white matter (i.e., Q1) to deeper white matter (i.e., Q4). To reiterate, Q1 corresponds to cerebellar fibers originating in the superficial layers of cerebellar cortex and includes parallel fibers. Additionally, Q2 corresponds to the portion of the tract that converges on the DCN, while Q3 is the point of divergence away from the DCN, corresponding to the more inferior portion of the superior cerebellar peduncles. Finally, Q4 corresponds to the more superior portion of the superior cerebellar peduncle that innervates the ventral tegmental area. Note that the quarter-wise analysis was performed in order to assess which subsection of the tract under study was most responsible for driving the brain-self-report effects.

**Table 3.** Correlations with Achenbach Self-Report (ASR) subscales and Fractional Anisotropy (FA) values for the right-to-right travelling polysynaptic tracts with the highest densities. A double dissociation between spinocerebellum (i.e., vermis and paravermis sending projections through the fastigial and interposed nuclei, respectively) and cerebrocerebellum (i.e., lateral cortical regions sending projections through the dentate) on somaticizing symptoms and social withdrawal, respectively. The N for all correlations is 101, and all confidence intervals are bootstrapped using 10,000 iterations. False Discovery Rate (FDR) correction for multiple comparisons was performed for the number of tracts within the polysynaptic (4 tracts) and monosynaptic (3 tracts; see Supplement) correlation domains for overall tract averages, separately. We used the FDR method to control for multiple comparisons, as we anticipated the effects to be smaller than those in the anatomy analyses and did not want to be unduly strict. Moreover, we chose to look at self-report measures for which we had *a priori* hypotheses, further warranting a less stringent approach to significance testing. Bootstrapped confidence intervals to further check for robustness of effects.

| Fractional Anisotropy Correlations: ASR Anxiety & Depression |  |  |  |  |
| --- | --- | --- | --- | --- |
| Segment/DCN | Section | Rho ($\rho$); $p$-value | $p$-value FDR | 95% CI:<br>Lower-Upper |
| a:vermis | **Avg.** | **.28; $p=.004$ \*\*** | **.016 \*\*** | **(.09, .46)** |
| | *Q-1* | .23; $p=.023$ * | | |
| | *Q-2* | .24; $p=.017$ * | | |
| | *Q-3* | .24; $p=.014$ * | | |
| | *Q-4* | .09; $p=.362$ ns | | |
| b:paravermis | **Avg.** | **.23; $p=.020$ \*** | **.032 \*** | **(.03, .42)** |
| | *Q-1* | .12; $p=.235$ ns | | |
| | *Q-2* | .21; $p=.034$ * | | |
| | *Q-3* | .23; $p=.020$ * | | |
| | *Q-4* | .09; $p=.391$ ns | | |
| c<sub>1</sub>:cerebro-cerebellum | **Avg.** | **.18; $p=.074$ †** | **.074 †** | **(-.02, .37)** |
| | *Q-1* | .05; $p=.647$ ns | | |
| | *Q-2* | .06; $p=.545$ ns | | |
| | *Q-3* | .20; $p=.044$ * | | |
| | *Q-4* | .09; $p=.379$ ns | | |
|  | **Avg.** | **.22; $p=.024$ \*** | **.032 \*** | **(.03, .40)** |
| | *Q-1* | .23; $p=.022$ * | | |
| | *Q-2* | .11; $p=.260$ ns | | |
| | *Q-3* | .22; $p=.026$ * | | |
| | *Q-4* | .06; $p=.523$ ns | | |
| Fractional Anisotropy Correlations: ASR Internalizing Symptoms |  |  |  |  |

| Segment/DCN | Section | Rho ($\rho$); $p$-value | | $p$-value FDR | 95% CI:<br>Lower-Upper |
| --- | --- | --- | --- | --- | --- |
| a:vermis | Avg. | .28; $p$=.004 | ** | .013 * | (.08, .47) |
| | $Q$-1 | .26; $p$=.010 | * | | |
| | $Q$-2 | .26; $p$=.009 | ** | | |
| | $Q$-3 | .23; $p$=.020 | * | | |
| | $Q$-4 | .06; $p$=.541 | ns | | |
| b:paravermis | Avg. | .26; $p$=.010 | * | .013 * | (.05, .44) |
| | $Q$-1 | .19; $p$=.058 | † | | |
| | $Q$-2 | .24; $p$=.018 | * | | |
| | $Q$-3 | .24; $p$=.014 | * | | |
| | $Q$-4 | .07; $p$=.462 | ns | | |
| c<sub>1</sub>:cerebro-cerebellum | Avg. | .21; $p$=.040 | * | .040 * | (.01, .39) |
| | $Q$-1 | .11; $p$=.279 | ns | | |
| | $Q$-2 | .12; $p$=.247 | ns | | |
| | $Q$-3 | .20; $p$=.041 | * | | |
| | $Q$-4 | .07; $p$=.470 | ns | | |
| | Avg. | .26; $p$=.010 | * | .013 * | (.06, .43) |
| | $Q$-1 | .27; $p$=.007 | * | | |
| | $Q$-2 | .20; $p$=.046 | * | | |
| | $Q$-3 | .24; $p$=.014 | * | | |
| | $Q$-4 | .06; $p$=.561 | ns | | |
| Fractional Anisotropy Correlations: ASR Social Withdrawal |  |  |  |  |  |
| Segment/DCN | Section | Rho ($\rho$); $p$-value | | $p$-value FDR | 95% CI:<br>Lower-Upper |
| a:vermis | Avg. | .15; $p$=.142 | ns | .142 ns | (-.04, .33) |
| | $Q$-1 | .25; $p$=.011 | * | | |
| | $Q$-2 | .15; $p$=.137 | ns | | |
| | $Q$-3 | .07; $p$=.484 | ns | | |
| | $Q$-4 | -.02; $p$=.819 | ns | | |
| b:paravermis | Avg. | .19; $p$=.064 | † | .085 † | (-.02, .38) |
| | $Q$-1 | .27; $p$=.007 | ** | | |
| | $Q$-2 | .21; $p$=.040 | * | | |
| | $Q$-3 | .16; $p$=.120 | ns | | |
| | $Q$-4 | .01; $p$=.908 | ns | | |
| c<sub>1</sub>:cerebro-cerebellum | Avg. | .20; $p$=.041 | * | .082 † | (.01, .39) |
| | $Q$-1 | .20; $p$=.046 | * | | |
| | $Q$-2 | .17; $p$=.085 | † | | |
| | $Q$-3 | .17; $p$=.091 | † | | |
| | $Q$-4 | .04; $p$=.663 | ns | | |
| | Avg. | .23; $p$=.019 | * | .076 † | (.04, .41) |
| | $Q$-1 | .21; $p$=.032 | * | | |

|  | *Q-2* | .30; p=.003 | ** |  |  |
| --- | --- | --- | --- | --- | --- |
|  | *Q-3* | .21; p=.033 | * |  |  |
|  | *Q-4* | .03; p=.737 | ns |  |  |
| Fractional Anisotropy Correlations: ASR Somaticizing Symptoms |  |  |  |  |  |
| Segment/DCN | Section | Rho ($\rho$); *p*-value | | *p*-value FDR | 95% CI:<br>Lower-Upper |
| a:vermis | **Avg.** | **.28; p=.004** | **\*\*** | **.014 \*** | **(.08, .48)** |
|  | *Q-1* | .29; p=.003 | ** |  |  |
|  | *Q-2* | .33; p=< .001 | *** |  |  |
|  | *Q-3* | .23; p=.023 | * |  |  |
|  | *Q-4* | .02; p=.878 | ns |  |  |
| b:paravermis | **Avg.** | **.27; p=.007** | **\*\*** | **.014 \*** | **(.07, .45)** |
|  | *Q-1* | .22; p=.027 | * |  |  |
|  | *Q-2* | .26; p=.010 | ** |  |  |
|  | *Q-3* | .24; p=.016 | * |  |  |
|  | *Q-4* | .03; p=.805 | ns |  |  |
| c<sub>1</sub>:cerebro-cerebellum | **Avg.** | **.19; p=.060** | **†** | **.060 †** | **(-.02, .38)** |
|  | *Q-1* | .16; p=.110 | ns |  |  |
|  | *Q-2* | -.08; p=.199 | ns |  |  |
|  | *Q-3* | .17; p=.090 | † |  |  |
|  | *Q-4* | .02; p=.830 | ns |  |  |
|  | **Avg.** | **.23; p=.020** | **\*** | **.027 \*** | **(.05, .41)** |
|  | *Q-1* | .27; p=.006 | ** |  |  |
|  | *Q-2* | .22; p=.031 | * |  |  |
|  | *Q-3* | .21; p=.032 | * |  |  |
|  | *Q-4* | .04; p=.702 | ns |  |  |
| Fractional Anisotropy Correlations: ASR Attention Problems |  |  |  |  |  |
| Segment/DCN | Section | Rho ($\rho$); *p*-value | | *p*-value FDR | 95% CI:<br>Lower-Upper |
| a:vermis | **Avg.** | **.20; p=.041** | **\*** | **.050 †** | **(.01, .39)** |
|  | *Q-1* | .10; p=.317 | ns |  |  |
|  | *Q-2* | .25; p=.014 | * |  |  |
|  | *Q-3* | .15; p=.135 | ns |  |  |
|  | *Q-4* | .03; p=.784 | ns |  |  |
| b:paravermis | **Avg.** | **.20; p=.050** | **†** | **.050 †** | **(-.004, .37)** |
|  | *Q-1* | .12; p=.247 | ns |  |  |
|  | *Q-2* | .18; p=.079 | † |  |  |
|  | *Q-3* | .21; p=.038 | * |  |  |
|  | *Q-4* | .06; p=.536 | ns |  |  |
| c<sub>1</sub>:cerebro-cerebellum | **Avg.** | **.20; p=.050** | **†** | **.050 †** | **(.002, .38)** |
|  | *Q-1* | .16; p=.108 | ns |  |  |
|  | *Q-2* | .12; p=.221 | ns |  |  |
| | *Q-3* | .18; $p=.081$ | † | | |
| | *Q-4* | .06; $p=.567$ | ns | | |
|  | **Avg.** | **.23; $p=.020$** | **\*** | **.050 †** | **(.05, .40)** |
| | *Q-1* | .29; $p=.003$ | ** | | |
| | *Q-2* | .20; $p=.042$ | * | | |
| | *Q-3* | .20; $p=.044$ | * | | |
| | *Q-4* | .06; $p=.540$ | ns | | |

† $p < .10$ , \* $p < .05$ , \*\* $p < .01$ , \*\*\* $p < .001$

Due to the large amount of data, statistics are reported only in the aforementioned tables, but are summarized in narrative form below. We note that it may be of interest to some researchers to know how the cerebello-VTA tract originating in the left cerebellum contributes to social and affective functioning—a matter that is not discussed at length in the current study (see **Table S12**). Due to the issue of multiple comparisons, we restricted the brain-self-report analyses to the right in an effort reduce the proliferation of analyses. However, we note that upon cursory investigation of left-to-left cerebello-VTA microstructure, only revealed significant relations between overall average FA in segment c_1_ crus II and ASR somatic complaints and attention problems. While other trends emerged, these results were not as compelling as those in the right-to-right tract, whose investigation is compelled by a data-driven anatomical approach.

#### 2.2.1. Correlations with Achenbach Adult Self Report

The findings for all ASR measures with FA across all 100 nodes of each polysynaptic tracts under study are summarized in **Figure 3**, while the findings for the monosynaptic analytic counterparts are summarized in supplemental **Figure S2**.

**Figure 3.**
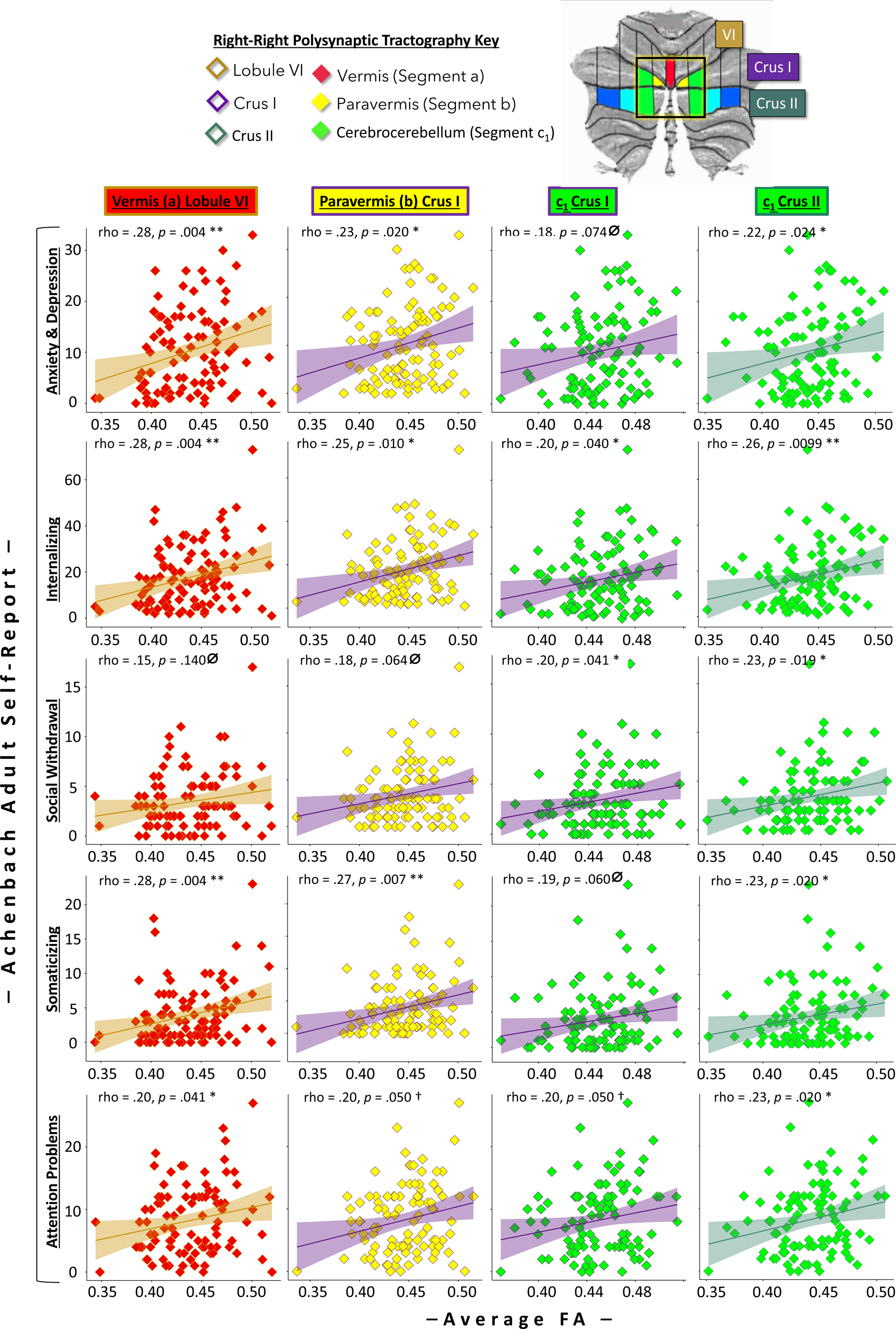
Summary of correlations between subscales on the ASR and FA averaged across all 100 nodes of the right-to-right travelling polysynaptic tracts of highest streamline density. Note that the *p*-values reflected here are not controlled for multiple comparisons. **Significance Flags:** l1 *p* > .05, † *p* =.05, * *p* < .05, ** *p* < .01

In line with the growing emphasis on dimensional, rather than categorical, conceptualizations of psychopathology (e.g., Snyder et al. (2023)), we not only examined anxiety and depression, but also associations between tractography and a broad measure of internalizing symptoms. In this way, we sought to apply a Research Domain Criteria (RDoC) approach to assess levels of negative affective symptomology across multiple dimensions (Cuthbert & Insel, 2013). For ease of reading, we limit our discuss to findings from the polysynaptic tract. Findings for the monosynaptic tract are discussed in narrative form at the end of the **Supplement.** Finally, the associated statistics are reported in supplemental **Table S13**.

##### Anxiety and Depression

Hypothesized correlations between the cerebello-VTA areas of highest tract density were identified between the vermal, paravermal and cerebrocerebellar portions of the tract originating in Crus II and anxiety and depression. All findings held when controlling for multiple comparisons. Quarter-wise analyses for polysynaptic tractographies revealed that Q3 emerged most often as driving this effect for each tract, followed by Q1 and Q2. Q4 never emerged as significant.

##### Internalizing Symptoms

Relations emerged between every polysynaptic tractography and internalizing symptoms, all of which survived FDR correction. The most implicated quarter was Q3, followed by Q2, and finally Q1. Q4 never emerged as significant.

##### Social Withdrawal

When averaging across all 100 nodes, the only relations to emerge with social withdrawal were cerebrocerebellar tracts originating in both crus I/II. Although these findings did not survive FDR correction, bootstrapped confidence intervals converged on significance for these tracts. The most implicated quarter was Q1, followed by Q2, and finally Q3. Q4 never emerged as significant.

##### Somatic Complaints

Somatic symptoms include a range of physical symptoms such as pain, dizziness, or shortness of breath. Somatic symptoms are distinct from but related to anxiety and depressive symptoms (Kong et al., 2022). Robust correlations between spinocerebellar segments of the cerebello-VTA tract and somatic complaints emerged. For the cerebrocerebellar portions of the tract, only the portion originating in crus II showed a significant relation. All findings survived multiple comparisons corrections, and all significant tracts implicated Q1, Q2, and Q3 equally. Q4 never emerged as significant.

##### Attention Problems

ASR attention problems were associated with lobule VI vermis-originating and crus II c_1_-originating FA. Although these associations did not hold after FDR correction, (p_FDR_ = .050 † in both cases), neither of the bootstrapped confidence intervals for these associations contained zero, bolstering the effects’ robustness. In terms of which portion of the tracts drove this effect, Q1-3 are implicated. It is important to note that Q2-3 are significant an equal number of times, while Q1 arises as significant only once.

#### 2.2.2. Correlations with NEO Five Factor Inventory

Correlations between whole tract FA for the spino- and cerebrocerebellar tract segments are reported in the complete correlation matrix in supplemental Table S11. As expected, there was no relation between cerebellum-VTA FA and the personality traits of conscientiousness and extroversion. However, we did find associations between FA and neuroticism and agreeableness. Hence, these relations are expounded upon in further detail below.

Unlike the analyses above, subjects were not matched on the NEO-FFI by biological sex. Consequently, we identified sex differences in neuroticism, with females exhibiting higher levels of each trait (W = 1592, p = .031 *, rrb = .25, 95% CI [.03, .45] and a trend in agreeableness W = 1542, p = .069 †, rrb = .21, 95% CI [-.01, .41]). To account for this in the following analyses, we report simple bivariate Spearman correlations and partial Spearman correlations accounting for sex. In each case, it is noted how controlling for sex changed the results.

##### Neuroticism

All polysynaptic tractographies of highest tract density correlated with NEO-FFI neuroticism, and all correlations survived FDR correction. Q3 was the most implicated, followed by Q2, while neither Q1 nor 4 were ever implicated. When controlling for sex, the correlations with overall tract FA held and survived FDR correction. However, the only remaining quarters implicated are Q3 and Q2, each to a lesser extent than previously.

##### Agreeableness

Lobule VI vermis-originating FA significantly correlated with Agreeableness. This finding did not survive FDR correction, though the bootstrapped confidence interval suggested a degree of robustness in the effect, as it did not contain 0. Though the vermal tract was the only one overall to reach significance, Q3 was significantly related to Agreeableness in all 4 polysynaptic tracts. These findings held when controlling for sex. The only change was that the relation between Q3 FA and Agreeableness for the Crus II c_1_ tract became a trend.

## 3. Discussion

### 3.1. Current findings

We mapped the structural connectivity between the cerebellum and VTA for the first time in humans. Using diffusion weighted imaging and probabilistic tractography, we found robust structural connectivity between both cerebellar cortex and the VTA, as well as cerebellar nuclei and the VTA. We then examined these connections at a higher level of granularity which revealed the following three findings. First, connections are most robust in the ipsilateral direction, although contralateral connections do exist. Second, there is higher tract density from the right cerebellum/right DCN to the VTA as compared to the left cerebellum/left DCN, suggesting some lateralization of function. Third, when we segmented crus I, II, and lobule IV, we found that there was a medial to lateral gradient such that the highest connectivity to the VTA was from the paravermis (segment b), with decreasing tract density as you moved lateral. Likewise, when we examined the monosynaptic tracts from the DCN to the VTA, the greatest density was from the interposed, followed by the fastigial, then the dentate nucleus.

We also charted the relationship between fractional anisotropy (FA) in areas of highest macrostructural density and *a priori* indices of socio-affective functioning. Further, we investigated whether associations between tract FA and self-report indices were driven by the full tract or by specific regions within the tract. We took at RDoC approach and assessed associations between tractography and broad measures of internalizing symptoms and negative affective symptomology. This revealed robust associations between FA in high-density portions of the cerebello-VTA tract and measures of psychiatric and personality self-report (see Figure 3). Strikingly, we identified that the cerebrocerebellar portion of the polysynaptic tract was uniquely associated with social withdrawal, while the monosynaptic fastigial and interposed tracts were uniquely associated with somatic complaints. Finally, we identified that the tract portion exiting the deep cerebellar nuclei was most responsible for driving the observed brain-self-report effects (e.g. Q3), followed by Q2, and then Q1, representing the more superficial (i.e., closer to the cerebellar cortical surface) partitions of the cerebello-VTA tracts.

### 3.2. The deep cerebellar nuclei, their inputs, and their outputs

The fastigial (medial) nucleus is the smallest of the DCN. Just lateral to it is the interposed nucleus, and lateral to that is the largest nucleus, the dentate (or lateral) nucleus (Schmahmann, 2019). In our study, we found the highest density of streamlines from the cerebellum to the VTA was from the paravermis, and predictably, from the interposed nucleus of the DCN. Prior histology findings have reported that these pathways project to the red nucleus, ventral lateral nucleus of the thalamus, periaqueductal grey, the superior colliculus (Nieuwenhuys, Voogd, & Huijzen, 2008), and the VTA (Novello et al., 2022). While the paravermis is known to calibrate motor control of the distal limbs (Unverdi & Alsayouri, 2023), other studies are slowly expanding the role of this region, beyond motor calibration. Paravermal lesions in mice result in downstream spikes in dopamine D1 receptor (DRD1) activity in the contralateral medial striatum, and dopamine transporter (DAT) attenuation in the dorsolateral striatum (Delis, Mitsacos, & Giompres, 2013). The influence of the interposed nucleus and the paravermis on dopamine activity is echoed in a study by S. R. Snider and Snider (1979), who found that paravermal lesions exert chronic effects on forebrain dopamine concentrations. This is interesting in light of our current findings due to the fact that DRD1 activity is implicated in a number of psychiatric conditions, as well as in social interest (Homberg et al., 2016). Furthermore, rodent tracer studies have revealed that most of the projections from the cerebellum to the nucleus accumbens (NAc) by way of the VTA originate in the interposed nucleus and travel ipsilaterally from the cerebellum (Oñate, Vera, Khatami, & Khodakhah, 2023), corroborating our anatomical findings.

The second highest density of streamlines was from the fastigial nucleus. The fastigial, by virtue of it being the output nucleus for the cerebellar vermis, has long been associated with emotion and motivation. Studies in rodents have found connections between the fastigial and periaqueductal gray, amygdala, septal nuclei, and hypothalamus (Cao et al., 2013; Fujita, Kodama, & du Lac, 2020; Heath & Harper, 1974; R. S. Snider & Maiti, 1976; Zhang, Wang, & Zhu, 2016). Lesions in the rat cerebellar vermis cause decreased innate fear, seen by an increased approach to natural predators (Supple, Leaton, & Fanselow, 1987).

In the current study, the most robust VTA-projecting region of the vermis came from lobule VI. Functional imaging studies of the cerebellum suggest several related possibilities for lobule VI of the vermis: that it is involved in saccadic eye movements, motor planning, bilateral hand movements, divided attention, visual working memory, and reconciliation of conflicting input (King, Hernandez-Castillo, Poldrack, Ivry, & Diedrichsen, 2019). Given the highly motoric nature of vermal lobule VI, it is possible that this portion of the cerebello-VTA tract may coordinate motivated seeking, or goal-directed behavior – the intersection of movement and reward – by way of the fastigial nucleus. This idea is bolstered by recent rodent work showing that GABAergic neurons in the VTA are involved in head movements in the appetitive pursuit of a moving target (Hughes et al., 2019; Masullo & Tripodi, 2019).

The least robust cerebello-VTA connections in our study were from the largest nucleus in the cerebellum, the dentate nucleus. The ventral dentate consists of projections to association cortices involved in cognitive and sensory processes (Bernard et al., 2014; Dimitrova et al., 2006; Dum & Strick, 2003; Palesi et al., 2015; Steele et al., 2017; Thurling et al., 2012), (Palesi et al., 2021). Relevant to the work discussed here, Bauer, Kerr, and Swain (2011) performed dentate nuclei lesions in rats and found that they exhibited decreased hedonic motivation (also see Low et al. (2021)). Another study showed that deactivation of VTA-projecting dentate neurons attenuated depression symptoms while excitation of this circuit exacerbated depression-like symptoms in stressed mice (Baek et al., 2022).

### 3.3. Relationship to preclinical work

Our findings are highly consistent with work from non-human animals. Findings from non-human animals show that cerebello-VTA connectivity exists that is both ipsilateral and contralateral, from all three DCN (Novello et al., 2022), (Judd et al., 2021), (Perciavalle et al., 1989), (R. S. Snider et al., 1976). Our findings also show ipsilateral and contralateral tracts from all three DCN to the VTA.

We also identified robust correlations between spino- and cerebrocerebellar portions of the cerebello-VTA tract and measures of depression symptomology in a normative sample. Our findings echo that of the chemo- and optogenetic rodent literature, especially that by Baek et al. (2022), who found that manipulation of crus I neurons sending projections to the VTA could either ameliorate or exacerbate depression-like symptoms. The circuit described in this paper as underpinning depression-like symptoms is homologous to our c_1_ crus I tract, in which we identified associations with internalizing and social withdrawal. Though our other crus I-originating tract originated in the paravermis, sending projections through the interposed nucleus, we also found that the VTA-sending tract originating in this lobule was associated with anxiety and depression, internalizing symptoms, somatic complaints, and trait neuroticism, consolidating the relevance of this cerebellar region in modulating affective processes.

Our behavioral findings also echo those of Carta et al. (2019), who found that activity between the DCN and VTA was necessary for mice to show a social preference. Interestingly, we found relations between the portions of the cerebello-VTA tract originating in crus I/II and social withdrawal scores on the ASR, a measure that gives an approximation of social interest in humans. Moreover, we found correlations with the portion of this tract that originates in lobule VI of the vermis and trait agreeableness. While not measuring social interest *per se*, this construct is inherently social in nature and provides a metric for assessing pro-sociality, altruism, and cooperation, and pleasantness (Wilson, Schneider, Arnold, Bienias, & Bennett, 2007), serving as a potential proxy for adaptive social proclivities that are stable across time (Atherton, Sutin, Terracciano, & Robins, 2022).

### 3.4. Relations to neuropsychiatric disorders

Our findings offer structural brain bases for emerging evidence that implicates the cerebellum in a host of neuropsychiatric disorders. The incidence of unipolar and bipolar depression increase tremendously following cerebellar injury in older adulthood (Frazier et al., 2022) and other mood symptoms, such as flattened affect, uncontrollable laughing or crying, and aberrant physiological markers of emotion processing are reported (Annoni, Ptak, Caldara-Schnetzer, Khateb, & Pollermann, 2003). Other studies have volumetric changes in the cerebellum in various mood disorders (reviewed in Frazier et al. (2022)), although the computational and/or biological mechanisms explaining this relationship are poorly understood.

Diminished responsivity to rewards and deficits in goal-directed behaviors are important features of depression (Paulus, 2015; Rizvi, Lambert, & Kennedy, 2018) and there is evidence of blunted reward signaling in depressed individuals (Kumar et al., 2018; Kumar et al., 2008). Things that are rewarding evoke dopaminergic signals that encode information about the discrepancy between reward expectancies and reward outcomes, to facilitate goal-directed behavior (Schultz, 2016). Previous research using probabilistic tractography in humans has implicated VTA-striatum and VTA-hippocampus connectivity in reward-motivated behaviors and psychopathology (Elliott, D’Ardenne, Mukherjee, Schweitzer, & McClure, 2022; Elliott, D’Ardenne, Murty, Brewer, & McClure, 2022; MacNiven, Leong, & Knutson, 2020). We extend these findings by suggesting that medial portions of the cerebellum may help regulate these circuits.

Cerebellar output is largely inhibitory. We previously predicted that stronger connectivity or higher FA in a cerebello-VTA circuit should be associated with higher levels of depressed mood and related internalizing symptoms. As seen in **Figure 3**, this is what we found across all metrics and ROIs.

### 3.5. Emotional embodiment in the cerebellum?

We also identified a robust set of correlations between varying portions of the cerebello-VTA tract and somatic symptoms, as revealed by the correlations with the fastigial and interposed monosynaptic tracts. This finding may relate to how sensorimotor and affective capacities intersect in the cerebellum in general, and in its midline in particular. One example of such convergence may be seen when cardiovascular modulation is perturbed—a function that is known to be subserved in part by the vermis—lending to a well-established increase in the prevalence of comorbid mood disorders (Bassett, 2016) and anxiety (Trotman et al., 2019). In fact, vermal lesions have been shown to impair heart rate conditioning in laboratory manipulations of fear-conditioned bradycardia (i.e., the heartrate deceleration shown to follow the immediate onset of a threat) (Battaglia, Nazzi, & Thayer, 2023). In a similar vein, posture has been shown to exert an influence on mood, such that those who retain an upright stance reap benefits for their general disposition (Awad, Debatin, & Ziegler, 2021), and maintain greater feelings of self-esteem and positive emotionality in the face of experimental paradigms of social stress (Nair, Sagar, Sollers, Consedine, & Broadbent, 2015). This is interesting in the context of our findings between FA in nearly identical portions of the cerebello-VTA tract and somatic and attention problems, as there is a suggestion that attending to and interpreting the physiological response to environmental provocation may be necessary for the resultant negative affective appraisal (Dalgleish, Dunn, & Mobbs, 2009) (see also Damasio (1996); Dunn, Dalgleish, and Lawrence (2006); Nauta (1971)). In this way, the cerebellum’s role in emotion may be in the coupling of emotions to action tendencies and physiological feedback.

### 3.6. Hierarchy of function by sagittal cerebellar segment

There is a bias towards aberrant social processes in higher order, phylogenetically newer portions of the cerebellum, denoted by sagittal segment c_1_, which correspond to the most medial portion of the cerebrocerebellum, as distinct from the spinocerebellum. Notably, the crus I/II c_1_ tracts were the only cerebello-VTA segments associated with ASR social withdrawal. Although a double dissociation of function is not fully present in comparing these results to that of the polysynaptic tracts’ associations with somaticizing, there is a bias towards phylogenetically older spinocerebellum’s role in more visceral emotional embodiment, which is evinced by the fact that the monosynaptic tracts carrying fibers away from the fastigial and interposed nuclei are uniquely associated with ASR somatic complaints. Moreover, it is critical to note that the monosynaptic dentate tractography showed a trend in its relationship with social withdrawal, with the quarter resting closest to the dentate (i.e., Q1), driving the trend. There may be a clearer double dissociation in the monosynaptic tractographies’ relations with ASR social withdrawal and somaticizing due to the possibility that there may be unique functionalities of the cerebellar cortex identified as the highest-density polysynaptic contributors to cerebello-VTA anatomy. Moreover, it is critical to note that while this potential hierarchy of function was identified in the ASR measures, it was not found to generalize to NEO-FFI measures of Agreeableness, which carries starkly social implications.

### 3.7. Limitations

First, our investigation only sought to detail the direct connections between the cerebellum and the VTA. There may be important indirect connections, such as those that may stop first in the red nucleus, thalamus, or hypothalamus. However, animal and human histology have suggested the relevance of the direct connections investigated here (Baek et al., 2022; Carta et al., 2019).

Second, we relied on streamlines as the macrostructural index of interest. Streamlines are known to be confounded by length, curvature, and branching, and may not correspond to tract density per se (Hoffman et al., 2022; Jones, Knosche, & Turner, 2013; Zhang et al., 2016). The number of streamlines reconstructed using DWI methods may not correspond to the actual number of axonal projections comprising the tract in question. Finally, probabilistic tracking algorithms such as the one employed in this study can be prone to false positive reconstructions (Maier-Hein et al., 2017). However, studies have validated diffusion-weighted tracking methods against known anatomy and tracer methods (Dyrby et al., 2007; Radwan et al., 2023; Sheng et al., 2021), with more recent evidence suggesting that constrained spherical deconvolution (CSD) probabilistic tracking is more accurate than traditional diffusion tensor imaging approaches (Martinez-Heras, Grussu, Prados, Solana, & Llufriu, 2021). Additionally, it is critical to note that DWI offers the unique ability to chart white matter connectivity in living organisms (Chanraud, Zahr, Sullivan, & Pfefferbaum, 2010). We hope that future investigators corroborate our findings using *ex vivo* techniques which do not suffer from these limitations.

Third, while we implemented a zebrin-informed partitioning of the posterior cerebellum, the sagittal segments under study do not precisely correspond to true zebrin expression. Our voxel size limited the granularity to chart this with precision. At its core, the sagittal segmentation scheme implemented in the current analysis is best conceptualized as a means of disambiguating spino- versus cerebrocerebellar cortical areas that send projections to distinct output nuclei and may represent a distinct hierarchy of complex behavior. Considering these caveats, future histological work is critical to investigate how individual differences in zebrin-negative versus zebrin-positive stripes of the neocerebellum may result in different behavioral phenotypes in the socio-affective domain.

Finally, we tested a normative sample who by definition, has relatively truncated variance at higher ends of symptom scales than a clinical sample who would have more severe social and affective dysregulation. It is possible that the trending relations identified between microstructure in the left-seeded portion of this tract may rise to the level of significance in a sample with more severe symptomology.

### 3.8. Conclusions and future directions

As the field of cognitive neuroscience turns its attention to the cerebellum to understand the pathophysiology of various neuropsychiatric disorders, it will be necessary to know where in the human cerebellum to look for the most robust projections. The current report found that the vermis and paravermis, along with the most medial portions of the cerebrocerebellum are most likely to provide input to the VTA, and that these connections predominantly originate in the right cerebellum and travel ipsilaterally. This study also provides the first evidence that cerebrocerebellar contributions to the cerebello-VTA tract may uniquely subserve individual differences in social interest, while the spinocerebellar deep nuclei (i.e., the fastigial and interposed) may be more responsible for emotional embodiment by way of somaticizing. In this way, this study sets the stage for a more cohesive human brain mapping of the cerebello-dopaminergic system, and follow-up investigations of its behavioral correlates in clinical samples.

This work also provides a guide about where to target the cerebellum if one wanted to modulate mood or motivation. Transcranial Magnetic Stimulation (TMS) of the cerebellar midline has already shown promise for alleviating negative symptoms in psychosis (Brady et al., 2019; Pierce, 2023). Our study lays the anatomical and functional groundwork for understanding how the diverse anatomy of cerebellar connectivity can manifest in neuropsychiatric dysfunction, and points towards how targeting distinct portions of this anatomy may ameliorate motivational and affective sequelae.

## 4. Materials & Methods

### 4.1. Data Availability

White matter fiber orientation distributions, subject-level region of interest/exclusion masks, and tractography files may be furnished upon request. Raw MRI, demographic, certain behavioral/self-report data are restricted by the National Institute of Health and the HCP. Because of this access to restricted data must be requested by fulfilling out the HCP’s Electronic Restricted Access Application. Finally, since much of the data in the current study is accessible to the larger research community, subject IDs will be available through OSF as of the date of publication. However, data files containing macro- and microstructural indices for the tracts under study have been deposited at OSF and will be made publicly available as of the date of publication. Further, exemplar tractography scripts and statistical code have been deposited at OSF and will also be publicly available as of the date of publication. Any additional information required to reanalyze the data reported in this paper is available from the lead contact upon request.

### 4.2. Participants

All imaging data was acquired from the minimally preprocessed S900 release of the Human Connectome Project dataset. For in-depth details concerning data acquisition parameters, preprocessing, and quality control for this dataset, see Van Essen and colleagues (Glasser et al., 2013; Marcus et al., 2013; Marcus et al., 2011; Sotiropoulos et al., 2013; Ugurbil et al., 2013; Van Essen et al., 2013; Van Essen et al., 2012).

Our goal was to have a sample size of ∼100, evenly split between biological male and female participants, who varied in their raw Achenbach Adult Self-Report (ASR) depression and anxiety subscale scores. We did this to have sufficient variability on affectively relevant behavioral measures for planned follow-up investigations on individual differences in tract microstructure as it relates to emotion dysregulation. Candidate participants were eliminated from the dataset if they did not have diffusion-weighted scans or T1-weighted scans, were left-handed, or did not have complete ASR data. Once these factors were considered, an even number of males and females were chosen. The remaining participants were ranked according to their raw ASR depression and anxiety subscale scores. Men and women at the low, middle, and high end of the ASR subscale were randomly selected in a balanced manner until an equal number of men and women were acquired. This was done by over selecting for those who had high raw scores on this ASR subscale, for whom subject’s scores were uniquely high (i.e., those for whom no ties were present on the ASR depression and anxiety subscale). From there, equal numbers of those for whom ties were present were randomly selected to ensure an egalitarian distribution of depression and anxiety scores. In the end, the demographic makeup of our sample was 72% white, 14% black or African American, 9% Asian/Native American/Pacific Islander, 2% unknown or not reported, and 3% mixed race. The average age of our subjects was 28.56 with a standard deviation of 3.69. Finally, the median income for the sample was within the bracket of $40K-$49,999, a figure that was based on reporting from 99 of the 101 participants under study, as two subjects did not answer this question.

To combat potential sex differences based on affective self-report, males and females were matched on the ASR anxiety and depression subscale. However, in the case of our secondary self-report measure, the NEO-Five Factor Inventory (NEO-FFI), males and females could not be matched, and so follow up analyses were performed controlling for sex in cases where a significant brain-behavior relation was identified. It is important to note that the current study did not make hypotheses regarding how race, ethnicity, or socio-economic status may impact tract macro- or microstructure, or how these factors may contribute to underlying anatomy or brain-behavior effects. As such, the current analyses should be replicated in the future with a more diverse cohort and a demographics-driven research question to ascertain whether these factors have an impact on anatomy and function.

### 4.3. Diffusion-Weighted Imaging Methods

The cerebellar ROIs included the deep cerebellar nuclei (DCN)—the fastigial, interposed, and dentate nuclei (Purves, Augustine, & Fitzpatrick, 2001; Rajasekhar, 2022)—and the cerebellar lobules VI, crus I and II. The ROIs were extracted from the Montreal Neurological Institute (MNI) probabilistic atlas for cerebellar lobules and deep nuclei (Diedrichsen, Balsters, Flavell, Cussans, & Ramnani, 2009; Diedrichsen et al., 2011) (see also https://www.diedrichsenlab.org/imaging/propatlas.htm). To prevent overlapping voxel coverage between the ROIs, we thresholded at varying levels for neighboring ROIs (see Table S1).

Cerebellar cortex has many distinct functional regions within each lobule, so we increased the granularity of our analysis by creating zebrin-informed sagittal segments of lobules VI, crus I and crus II. This was performed after setting the minimum threshold for each at 25 using the *fslmaths* command in FSL. This resulted in left and right regions for each lobule that were 20-voxels and 25-voxels wide, respectively. Each left and right lobule was then partitioned into 5-voxel segments along the x-axis, resulting in 4 segments per hemisphere for lobule VI, and 5 segments per hemisphere for crus I and II. Sagittal divisions are labeled by number, with higher numbers indicating more lateral portions of each lobule. For the purposes of the current analysis, segment 1 of the cerebellar lobules has been assigned the title of paravermis, or “spinocerebellum”, due to its medial positioning immediately adjacent to the vermis. Thus, segment b sends projections to the interposed nucleus (IN) in subsequent analyses. All other segments (i.e., c_1_ – c_3_ for lobule VI, and c_1_ – c_4_ for crus I and II) are considered part of the cerebrocerebellum, and therefore project to the dentate nucleus (DN).

Since the vermis contains subdivisions corresponding to each cerebellar lobule, and the particular portions of this structure that may be most important for social and affective functioning is still uncharted, for the sake of the current analysis we only investigate portions of the vermis that correspond to the cerebellar lobules of interest. Note that all vermal projections synapse onto the fastigial nucleus (FN) before leaving the cerebellum.

Brainstem ROIs were extracted from the brainstem navigator atlas, derived from 7 Tesla (T) *in vivo* Magnetic Resonance Imaging (MRI) using multiple contrasts and segmentation methods (https://www.nitrc.org/projects/brainstemnavig/) (Bianciardi et al., 2016). We used the bilateral VTA from this atlas as our target ROI, and further used the red nucleus and Inferior Colliculus (IC) as exclusion masks in the tractography analyses described below.

To delineate these transformation parameters, a series of linear registrations were performed. First, subjects’ T1-weighted images were skull-stripped using FSL’s brain extraction tool, which executed a bias field and neck cleanup. Next, subjects’ topup-corrected, time-collapsed b0 images were registered to their respective T1-weighted anatomical images with six degrees of freedom and a correlation ratio cost function, yielding a diffusion to structural space conversion matrix. The same registration parameters were used to warp subjects’ T1-weighted images to the 2 x 2 x 2mm^3^ MNI space brain template, creating a structural to standard space conversion file. Next, the inverse of the two aforementioned matrices was taken to produce a structural to diffusion space and a standard to structural space conversion matrix, respectively. Last, to acquire the diffusion to standard space transformation parameters, the structural to standard space and diffusion to structural space matrices were concatenated. Finally, the inverse of this file was taken to create the standard to diffusion space conversion matrix that was used to convert the spheres to subject space. This procedure was enacted to avoid performing transformations directly on images that transcend two different imaging modalities (i.e., T1-weighted and diffusion MRI). Finally, the MNI to diffusion space transformation matrix was used in conjunction with FSL’s non-linear registration tool, *fnirt*, to warp ROIs to diffusion space (Andersson, Jenkinson, & Smith, 2010).

Further, it is critical to note that one key aim of the current investigation was to reveal monosynaptic, direct connections between the cerebellum and VTA. This is challenging due to the proximity of the VTA to the red nucleus and thalamus, which are waypoints in the cerebello-thalamo-cortical tract (CTC)—the largest cerebellar output tract (Palesi et al., 2015). Therefore, to dissociate the cerebello-VTA pathway from the CTC, we excluded the thalamus and red nucleus bilaterally. To prevent overlap with the VTA, the VTA was subtracted from the red nucleus once in subject space, and the difference was used in the exclusion mask.

To ensure extraneous fibers were not reconstructed as part of this investigation, we also excluded the brainstem inferior to, and including the pons, the cerebellum contralateral to where each tractography was seeded, the VTA contralateral to that being targeted, and all lobes of the cerebral cortex. In the case of the decussating tract, we forced decussation at the level of the IC via the Wernekink commissure. To do this, we identified the last slice of the IC ROIs from the Brainstem Navigator atlas in MNI space and excluded all voxels superior to that slice in the hemisphere ipsilateral to the cerebellar seed region.

MRTrix3 (https://www.mrtrix.org/) was used to estimate voxel-wise response functions for each principal neural tissue compartment, including grey matter, white matter, and cerebral spinal fluid, using the *dhollander* algorithm (Dhollander, Mito, Raffelt, & Connelly, 2019; Dhollander, Raffelt, & Connelly, 2016, 2018). We fit multi-tissue orientation distribution functions for each macroscopic tissue type using the *msmt_csd* argument in the *dwi2fod* command. This approach capitalized on the distinct diffusion properties of different tissue types measured by the multi-b-value, High Angular Resolution Diffusion Imaging (HARDI) sequence in the HCP (Jeurissen, Tournier, Dhollander, Connelly, & Sijbers, 2014).

Probabilistic tractography was performed using the iFOD2 algorithm in MRTrix3 using the *tckgen* command. Note that for all tractographies, streamline selection criteria were disabled. Further, all tractographies seeded in the cerebellar cortex were performed unidirectionally, with the VTA designated as both an inclusion and stopping criteria. To make conjectures about tract density, we seeded a fixed number of streamlines per voxel in the seed mask, with random placement of the seeds throughout. To control for differing seed size of distinct cerebellar cortical areas, we determined the lowest volumetric occurrence of any of the cortical seed regions within the cohort. We found the smallest size in any subject to be that of the vermis corresponding to right crus II, with a volume of 329 voxels. Using 5,000 seeding attempts per voxel as our default selection criteria, we calculated the total number of viable seeds for all lobule-based tractographies by multiplying 329 voxels by 5,000 seed attempts per voxels, resulting in 1,645,000 total seeds for each tractography. The seeding-attempts per voxel for all other lobule-based tractographies was anchored by the smallest volumetric occurrence for each seed in the cohort and was calculated by dividing the smallest seed volume for each tractography in our sample by 1,645,000 total seeds. This information is summarized in table S2.

For tractographies seeded in the DCN, tracking was performed both unidirectionally and bidirectionally. The latter was done to evaluate where each nucleus receives input from the cerebellar cortex on the way to the ventral tegmental area. Moreover, due to the small size of the DCN relative to the portions of cerebellar cortex under study, the total number of seeds was anchored at the smallest occurrence of any of the DCN. In this case, the smallest nucleus was found in a subject whose right fastigial nucleus was 91 voxels. Therefore, for all DCN tractographies, the total number of seeds was set at 91 voxels * 5,000 seeding attempts per voxel, yielding a total seed count of 455,000 for each tractography. The value of seeds per voxel for all other DCN-seeded tractographies was determined by dividing the smallest occurrence of each DCN in the cohort, divided by the total number of seeds (455,000). This information is summarized in table S2. Subject-wise streamlines and tract length were extracted using the MrVista package (https://github.com/vistalab/vistasoft) in MATLAB version 2017b (9.3).

Polysynaptic tracts (i.e., those originating in the cerebellar cortex, synapsing onto the DCN, and terminating in the VTA) from regions found to have the highest density of streamlines, as well as each monosynaptic tract originating in each DCN, were split into 100 nodes using Automated Fiber Quantification through Python (PyAFQ) (Kruper et al., 2021). That is, tractography between the cerebellum and VTA was funneled into PyAFQ, which split the tracts into 100 equidistant nodes. We then extracted the diffusion metric, Fractional Anisotropy (FA), at each node, for each tract, for every subject. Therefore, different “FA tract profiles” were output for each subject (Yeatman, Dougherty, Myall, Wandell, & Feldman, 2012). This is important as FA values are not the same along the whole tract, so in averaging across the entirety of the tract systematic variability across each bundle is diminished. Further, to get a total FA profile for each tract, FA was averaged across each of the 100 nodes. Then, to delve deeper into the portions of each tract that were found to correlate with self-report indices, we split the 100 nodes into four quarters by averaging FA every 25 nodes. In keeping with the data-driven approach and to reduce the number of multiple comparisons in the current analysis, FA in the right-seeded ipsilaterally travelling tracts were quantified, as these tracts were found to have the highest density of streamlines traveling to the VTA.

Based on known neuroanatomy, we can make some assumptions about each quarter. For the polysynaptic tracts, quarter one (Q1) likely corresponds to the granule cells, parallel fibers, and Purkinje cell fibers in the molecular and Purkinje cell layers. Quarter two (Q2) corresponds to the granular layer containing Purkinje cell axons and climbing fibers at the juncture where these projections hit the DCN. Quarter three (Q3) corresponds to the mossy fibers and DCN outputs converging on the superior cerebellar peduncles. Finally, quarter four (Q4) corresponds to the point of VTA innervation in the midbrain. The anatomical breakdown of the quarters for the polysynaptic tracts are exemplified in a representative tract in **Figure 4**.

**Figure 4.**
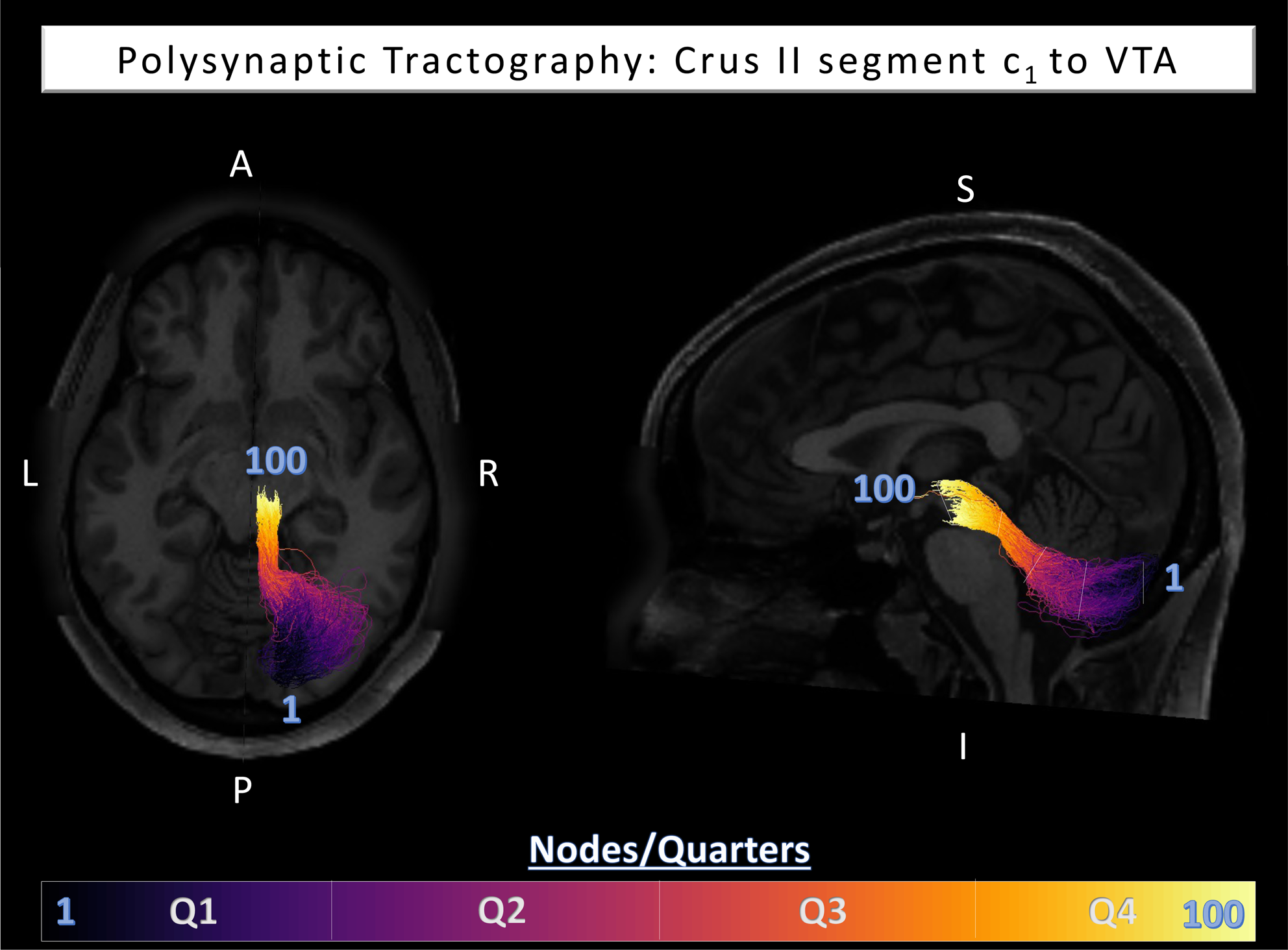
Visualization for a representative polysynaptic tractography originated in Crus II of segment c_1_ of the cerebrocerebellum. The number 1 indicates where the node-wise segmentation originated, while 100 represents the where the segmentation ended. The boundaries of each quarter are indicated by white lines on the sagittal view of the tract, and the color coding corresponding to each quarter is summarized in the color bar at the bottom of the figure. In cases of monosynaptic tractographies that were also split into quarters, this segmentation involves the equal quarterization of Q2 to Q4 visualized above.

For the monosynaptic tracts, Q1 is the point of departure from the DCN within the cerebellum. Q2 and Q3 reflect the inferior and superior portions of the superior cerebellar peduncles, respectively. Finally, Q4 is the point of VTA innervation in the midbrain.

### 4.4. Self-Report Measures

To assess associations between tract microstructure and psychiatric and life function in the domains of negative affect, anxiety, depression, and social support, we examined measures form the Achenbach Adult Self Report for Ages 18-59 (ASR; Achenbach, 2009). To explore specific domains of functioning, we examined ASR Syndrome Scales as quantified by the 123 items from Section VIII of the assessment. We predicted we would see correlations between the anxiety and depression, social withdrawal, and internalizing symptom subscales. In a set of exploratory investigations, we also assessed associations with somatic complaints and attention problems. Finally, we evaluated correlations between externalizing symptoms and aggressive behavior as control analyses. Since this is a normative sample, we probed for associations using raw scores instead of age and/or gender adjusted percentile scores to retain sufficient variability to detect effects.

Next, to examine relations between tract microstructure and socially and affectively relevant personality traits, we looked at correlations with the Neuroticism/Extroversion/Openness Five Factor Inventory (NEO-FFI), a shortened, 60-item version of the original five factor assessment (Costa & McRae). The FFI has exhibited strong reliability and validity cross culturally and is thought to reflect key components of human personality (Heine & Buchtel, 2009; McCrae & Costa, 2004). In the current analysis, we looked at scores on Neuroticism (N), Extroversion/Introversion (E), and Agreeableness (A) subscales as outcomes of interest, as these measures provide insights into stable measures of emotional (N) and social (E and A) functioning. As a control outcome, we examine correlations with trait Conscientiousness (C), as it is orthogonal to socio-affective disposition.

### 4.5. Statistical Analyses

All statistical analyses for the anatomical aims were performed in RStudio version 1.4.1717 (https://www.r-project.org/). For the brain-self-report analyses, statistics were performed in JASP version 0.18 (JASP Team, 2023), a graphical-user-interface-based software whose engine runs in R.

#### 4.5.1. Anatomical Analyses

Focused, pairwise contrasts were performed to ascertain the following comparisons pertaining to tract density: (1) contralaterally vs. ipsilaterally synapsing tracts, performed for each cerebellar segment; (2) right- versus left-traveling tracts; (3) vermal vs. paravermal, vermal vs. cerebrocerebellar, and paravermal vs. cerebrocerebellar tracts; (4) tracts originating in crus I vs. crus II, crus I vs. lobule VI, and crus II vs. lobule VI; (5) Projections to the VTA from the dentate nucleus vs. interposed nucleus, dentate nucleus vs. fastigial nucleus, and interposed nucleus vs. fastigial nucleus. All contrasts were performed using paired Wilcoxon signed-rank tests in the *rstatix* package. To get effect sizes, we used the *wilcox_effsize()* command. Moreover, we computed 95% percentile confidence intervals bootstrapped around the effect size using 10,000 iterations.

Note that the results for the polysynaptic circuitry are presented 3 different ways: (1) without any controls; (2) after controlling for size of the DCN ROI in each projection route (dividing number of streamlines for each tract by the volume of the relevant DCN ROI on a subject-wise basis); (3) after controlling for tract length (dividing the number of streamlines by tract length). In this way, DCN ROI volume and tract length are treated as potential confounds, and the extent to which the 3 results converge is assessed to ascertain the relative importance of each in relation to our findings. For a visualization of the tractographies performed in the current analysis, see **Figure 5**.

**Figure 5.**
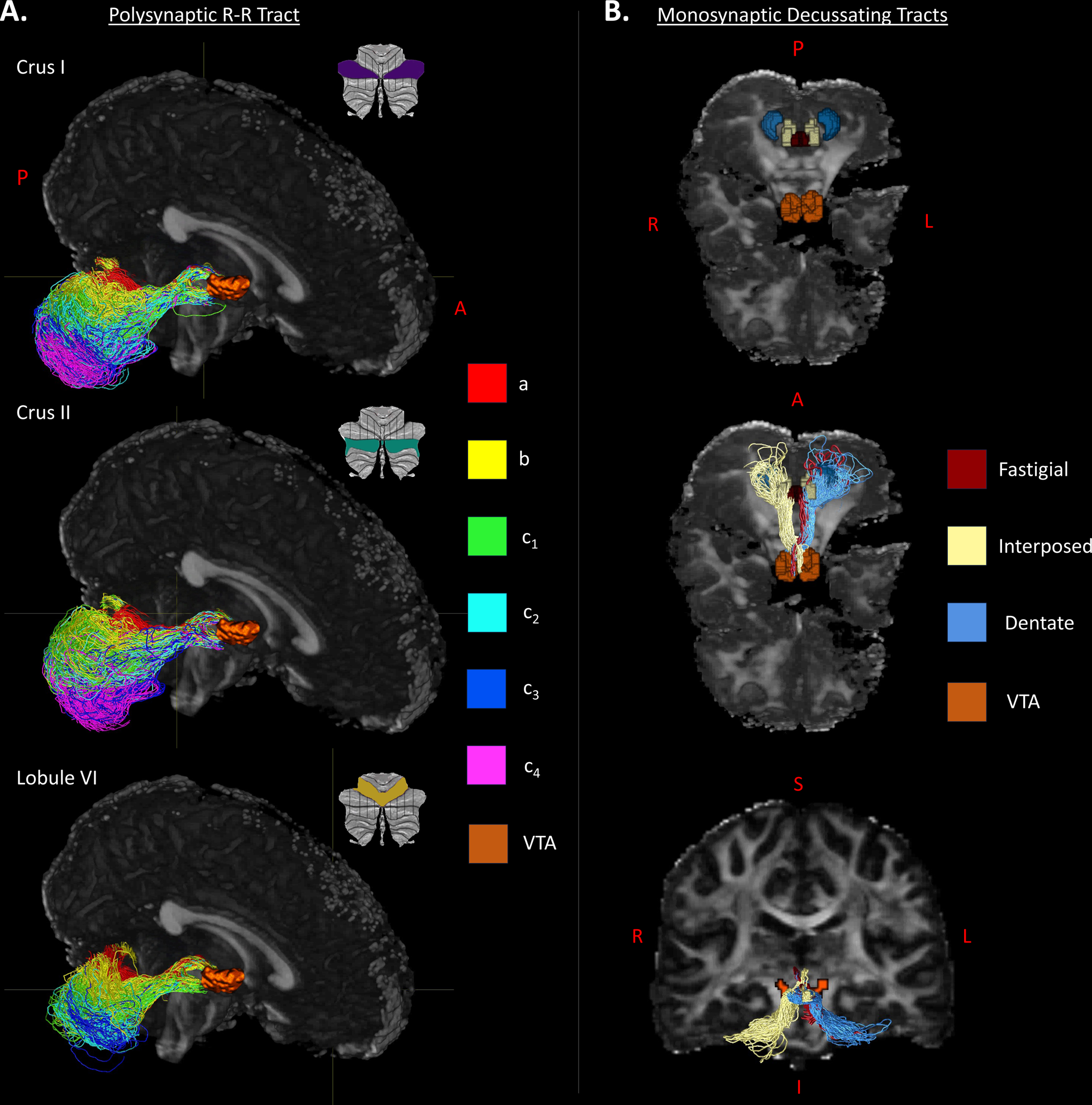
Tractography results in a representative subject of connections from cerebellum to deep cerebellar nuclei to VTA. A. Right hemisphere ipsilaterally-traveling polysynaptic tracts are shown for sagittal segments (a, b, and c_1_ – c_4_) within each cerebellar lobule. B. Anatomical Regions-Of-Interest (ROIs) include the 3 deep cerebellar nuclei (Fastigial, Interposed, and Dentate), and the VTA. Tractography results of the monosynaptic tracts between the deep cerebellar nuclei and VTA, color-coded by the nuclei from which they were seeded.

#### 4.5.2. Brain-Self-Report Analyses

Bivariate non-parametric Spearman correlations are reported for all analyses between tract segment FA and ASR and NEO-FFI personality traits. Follow-up exploratory quarter-wise tract correlations with behavior were also performed in a bivariate fashion. Moreover, the frequency of each quarter’s emergence as the driver of the overall brain-behavior effect in question was investigated. For correlations between tract microstructure and NEO-FFI measures on which subjects were not able to be matched by sex, partial correlations controlling for sex were performed. Since the groups were matched for sex on the ASR, and since age was not a confound in the case of the ASR or NEO-FFI, neither multivariate regressions, nor partial correlations were performed.

## Acknowledgements

We would like to thank Drs. Volker Coenen, Marco Reisert, and Sascha du Lac for their insights into cerebello-midbrain connectivity and comments on the cerebellar connections to each deep cerebellar nucleus, respectively. We would also like to thank Katie Jobson, Haroon Popal, and Vishnu Murty for contributing their knowledge of cerebellar and reward circuitry. Moreover, data were provided by the Human Connectome Project, WU-Minn Consortium (Principal Investigators: David Van Essen and Kamil Ugurbil; 1U54MH091657) funded by the 16 NIH Institutes and Centers that support the NIH Blueprint for Neuroscience Research; and by the McDonnell Center for Systems Neuroscience at Washington University.

