## Supplemental_Materials for "An *in vivo* Dissection, and Analysis of Socio-Affective Symptoms related to Cerebellum-Midbrain Reward Circuitry in Humans"

**Supplement**

**Table S1.** Thresholds for ROIs. These threshold values were fed into *fslmaths* using the *-thr* argument in order to yield the final ROI masks.

| Cerebellar Cortex | | | | | | |
| --- | --- | --- | --- | --- | --- | --- |
|  | Crus I | | Crus II | | Lobule VI | |
|  | Hemisphere | Vermis | Hemisphere | Vermis | Hemisphere | Vermis |
| Left | 25 | 1 | 25 | 20 | 25 | 20 |
| Right |  |  |  |  |  |  |
| Deep Cerebellar Nuclei | | | | | | |
|  | Fastigial Nucleus | | Interposed Nucleus | | Dentate Nucleus | |
| Left | 5 | | 25 | | 45 | |
| Right |  |  |  |  |  |  |

**Table S2.** ROI and tractography parameters. Anchor volume for each seed region is reported in voxels. Seeds per voxel were calculated by dividing the anchor volume in voxels by 1,645,000 in the case of the polysynaptic tractography, and by 455,000 in the case of the monosynaptic tractography. Anc = anchor; Vol=volume; vox = voxel;

| Polysynaptic Anchor & Tractography Parameters | | | | | | | | | | | | |
| --- | --- | --- | --- | --- | --- | --- | --- | --- | --- | --- | --- | --- |
| Crus I | | | | | | | | | | | | |
| Hemi-sphere | Segment | | | | | | | | | | | |
|  | a | | b | | c_1_ | | c_2_ | | c_3_ | | c_4_ | |
|  | Anc. Vol | Seeds/ vox | Anc. Vol | Seeds/ vox | Anc. Vol | Seeds/ vox | Anc. Vol | Seeds/ vox | Anc. Vol | Seeds/ vox | Anc. Vol | Seeds/vox |
| Left | 795 | 2069 | 870 | 1891 | 1437 | 1145 | 2337 | 704 | 3397 | 484 | 2346 | 701 |
| Right | 502 | 3277 | 898 | 1832 | 1522 | 1081 | 2449 | 672 | 3510 | 469 | 2349 | 700 |
| Crus II | | | | | | | | | | | | |
| Hemi-sphere | Segment | | | | | | | | | | | |
|  | a | | b | | c_1_ | | c_2_ | | c_3_ | | c_4_ | |
|  | Anc. Vol | Seeds/ vox | Anc. Vol | Seeds/ vox | Anc. Vol | Seeds/ vox | Anc. Vol | Seeds/ vox | Anc. Vol | Seeds/ vox | Anc. Vol | Seeds/vox |
| Left | 407 | 4042 | 1531 | 1074 | 1757 | 936 | 1689 | 974 | 2218 | 742 | 1725 | 954 |
| Right | 329 | 5000 | 1287 | 1278 | 1820 | 904 | 1692 | 972 | 1907 | 863 | 1686 | 976 |
| Lobule VI | | | | | | | | | | | | |
| Hemi-sphere | Segment | | | | | | | | | | | |
|  | a | | b | | c_1_ | | c_2_ | | c_3_ | | | |
|  | Anc. Vol | Seeds/ vox | Anc. Vol | Seeds/ vox | Anc. Vol | Seeds/ vox | Anc. Vol | Seeds/vox | Anc. Vol | | Seeds/vox | |
| Left | 1262 | 1303 | 1445 | 1138 | 2180 | 755 | 2589 | 635 | 1221 | | 1347 | |
| Right | 964 | 1706 | 1433 | 1148 | 2072 | 794 | 2451 | 671 | 1093 | | 1505 | |
| Monosynaptic Anchor & Tractography Parameters | | | | | | | | | | | | |
| Hemi-sphere | Deep Cerebellar Nuclei | | | | | | | | | | | |
|  | Fastigial Nucleus | | | | Interposed Nucleus | | | | Dentate Nucleus | | | |
|  | Anc. Vol | | Seeds/ vox | | Anc. Vol | | Seeds/ vox | | Anc. Vol | | Seeds/vox | |
| Left | 94 | | 4840 | | 203 | | 2241 | | 322 | | 1413 | |
| Right | 91 | | 5000 | | 199 | | 2286 | | 318 | | 1431 | |

**Table S3.** Success rate for each tractography by number of non-zero reconstructions.

| Polysynaptic Success Rate | | | | | | |
| --- | --- | --- | --- | --- | --- | --- |
| Crus I | | | | | | |
| Direction | Segment | | | | | |
|  | a | b | c_1_ | c_2_ | c_3_ | c_4_ |
| R-R | 100 | 101 | 101 | 101 | 101 | 98 |
| L-L | 97 | 101 | 98 | 98 | 98 | 91 |
| R-L | 100 | 101 | 100 | 99 | 96 | 93 |
| L-R | 83 | 91 | 94 | 83 | 78 | 64 |
| Crus II | | | | | | |
| Direction | Segment | | | | | |
|  | a | b | c_1_ | c_2_ | c_3_ | c_4_ |
| R-R | 101 | 101 | 101 | 101 | 101 | 99 |
| L-L | 101 | 100 | 99 | 99 | 94 | 93 |
| R-L | 100 | 101 | 98 | 100 | 98 | 90 |
| L-R | 90 | 94 | 90 | 88 | 78 | 73 |
| Lobule VI | | | | | | |
| Direction | Segment | | | | | |
|  | a | b | c_1_ | c_2_ | c_3_ | |
| R-R | 101 | 101 | 99 | 99 | 98 | |
| L-L | 101 | 101 | 95 | 92 | 89 | |
| R-L | 100 | 98 | 96 | 93 | 94 | |
| L-R | 98 | 91 | 78 | 66 | 53 | |
| Monosynaptic Success Rate | | | | | | |
| Direction | Deep Cerebellar Nuclei | | | | | |
|  | Fastigial Nucleus | | Interposed Nucleus | | Dentate Nucleus | |
| R-R | 101 | | 101 | | 101 | |
| L-L |  |  |  |  | 100 | |
| R-L |  |  |  |  | 101 | |
| L-R | 97 | | 99 | | 94 | |

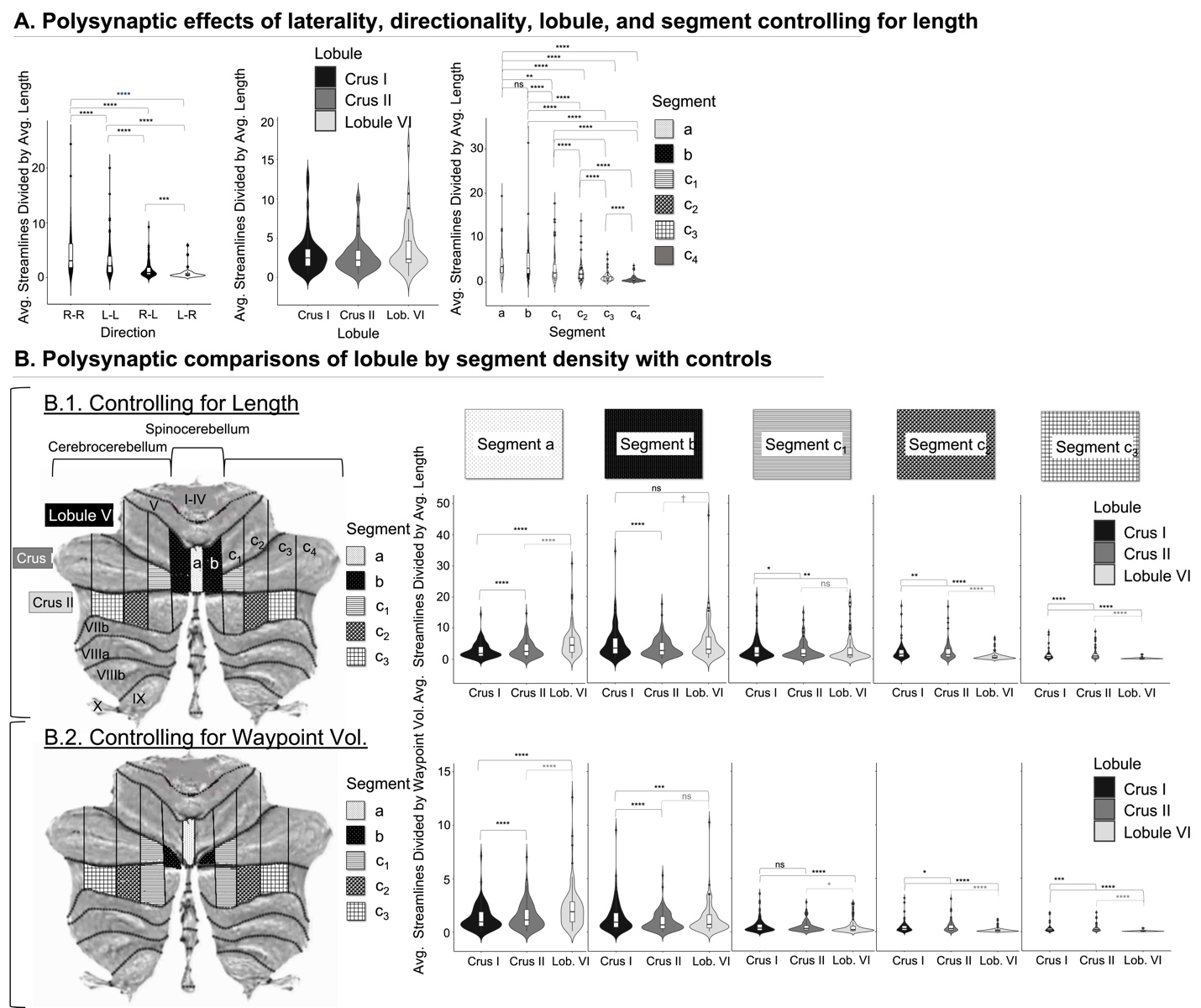

**Figure S1.** Illustrations of main effect and interaction effect statistics with controls not illustrated in **Figure 2.** Section (A) shows the statistical results of the main effects under study when controlling for average tract length. Section (B) shows the statical results of the segment by lobule interactions when controlling for average tract length (B.1.) and waypoint volume (B.2.). Note that the flat maps on the left in section (B) contain pattern-coded highlights of which lobule contained the greatest streamline density within each segment when taking into account different controls. Note that for the flat map in (B.1.), lobule VI and crus I are highlighted, as they both contained more connectivity than crus II, but did not differ from each other. Moreover, as in **Figure 2C**, no difference was found between crus I and II in segment c_4_, and so it remains both unhighlighted in the flap map and unrepresented in the violin plots.

**Table S4.** **(A)** Nonparametric pairwise comparisons using Wilcoxon sign-rank test investigating streamline count differences based on tract direction, lobule, and segment for polysynaptic tractographies; and **(B)** direction and deep nuclei macrostructural differences in monosynaptic tractographies.

| **A. Polysynaptic Tractography Main Effects: Controlling for Average Streamline Length** | | | | | | | | | | | |
| --- | --- | --- | --- | --- | --- | --- | --- | --- | --- | --- | --- |
| Direction | | | | | | | | | | | |
| Measure 1 | Measure 2 | *n* 1 | *n* 2 | W | *p* | *p_bonferroni_* | | *Eff. Size* | CI 95% | | Magnitude |
| R-R | L-L | 92 | 80 | 2377 | 2.25e-06 | 1.35e-05 | **** | .54 | .36 | .70 | large |
|  | R-L |  |  | 3128 | 3.95e-14 | 2.37e-13 |  | .85 | .82 | .87 |  |
|  | L-R |  | 32 | 496 | 9.31e-10 | 5.59e-09 |  | .87 | .87 | .87 |  |
| L-L | R-L | 80 | 80 | 2326 | 1.95e-09 | 1.17e-08 |  | .71 | .58 | .81 |  |
|  | L-R |  | 32 | 527 | 9.31e-10 | 5.59e-09 |  | .87 | .85 | .87 |  |
| R-L |  |  |  | 444 | 3.57e-05 | 2.14e-04 | *** | .69 | .46 | .84 |  |
| Lobule | | | | | | | | | | | |
| Measure 1 | Measure 2 | *n* 1 | *n* 2 | W | *p* | *p_bonferroni_* | | *Eff. Size* | CI 95% | | Magnitude |
| Crus I | Crus II | 49 | 56 | 507 | .328 | .984 | ns | .15 | .007 | .45 | small |
|  | Lobule VI |  | 43 | 267 | .441 | > .999 |  | .13 | .008 | .45 |  |
| Crus II |  | 56 |  | 269 | .323 | .969 |  | .17 | .008 | .48 |  |
| Segment | | | | | | | | | | | |
| Measure 1 | Measure 2 | *n* 1 | *n* 2 | W | *p* | *p_bonferroni_* | | *Eff. Size* | CI 95% | | Magnitude |
| a | b | 77 | 83 | 980 | 8.80e-02 | > .999 | ns | .20 | .02 | .42 | small |
|  | c_1_ |  | 75 | 1492 | 3.05e-04 | 5.00e-03 | ** | .46 | .23 | .67 | moderate |
|  | c_2_ |  | 62 | 1094 | 2.07e-07 | 3.10e-06 | **** | .68 | .49 | .83 | large |
|  | c_3_ |  | 48 | 690 | 1.28e-09 | 1.92e-08 |  | .84 | .75 | .87 |  |
|  | c_4_ |  | 56 | 945 | 4.55e-13 | 6.82e-12 |  | .87 | .86 | .87 |  |
| b | c_1_ | 83 | 75 | 2456 | 1.24e-09 | 1.86e-08 |  | .71 | .57 | .82 |  |
|  | c_2_ |  | 62 | 1716 | 3.64e-10 | 5.46e-09 |  | .82 | .74 | .86 |  |
|  | c_3_ |  | 48 | 1128 | 1.42e-14 | 2.13e-13 |  | .87 | .87 | .87 |  |
|  | c_4_ |  | 56 | 1431 | 2.46e-10 | 3.69e-09 |  |  |  |  |  |
| c_1_ | c_2_ | 75 | 62 | 1696 | 9.50e-10 | 1.42e-08 |  | .80 | .71 | .86 |  |
|  | c_3_ |  | 48 | 1128 | 1.42e-14 | 2.13e-13 |  | .87 | .87 | .87 |  |
|  | c_4_ |  | 56 | 1485 | 1.67e-10 | 2.50e-09 |  |  |  |  |  |
| c_2_ | c_3_ | 62 | 48 | 990 | 1.14e-13 | 1.71e-12 |  |  |  |  |  |
|  | c_4_ |  | 56 | 1225 | 3.55e-15 | 5.32e-14 |  |  |  |  |  |
| c_3_ |  | 48 |  | 946 | 2.27e-13 | 3.40e-12 |  |  |  |  |  |
| **B. Monosynaptic Tractography Main Effects: Controlling for Average Streamline Length** | | | | | | | | | | | |
| Direction | | | | | | | | | | | |
| Measure 1 | Measure 2 | *n* 1 | *n* 2 | W | *p* | *p_bonferroni_* | | *Eff. Size* | CI 95% | | Magnitude |
|  |  |  |  |  |  |  |  |  | Lower | Upper |  |
| R-R | L-L | 101 | 100 | 4098 | 6.42e-08 | 3.85e-07 | **** | .54 | .39 | .67 | large |
|  | R-L |  | 101 | 5108 | 9.70e-18 | 5.82e-17 |  | .85 | .83 | .87 |  |
|  | L-R |  | 93 | 4371 | 5.66e-17 | 3.40e-16 |  | .87 | .87 | .87 |  |
| L-L | R-L | 100 | 101 | 4614 | 6.92e-13 | 4.15e-12 |  | .62 | .62 | .80 |  |
|  | L-R |  | 93 | 4370 | 5.85e-17 | 3.51e-16 |  | .87 | .87 | .87 |  |
| R-L |  | 101 |  | 4081 | 3.85e-13 | 2.31e-12 |  | .65 | .65 | .83 |  |
| Deep Cerebellar Nuclei | | | | | | | | | | | |
| Measure 1 | Measure 2 | *n* 1 | *n* 2 | W | *p* | *p_bonferroni_* | | *Eff. Size* | CI 95% | | Magnitude |
|  |  |  |  |  |  |  |  |  | Lower | Upper |  |
| FN | IN | 97 | 99 | 962 | 3.62e-07 | 1.09e-06 | **** | .52 | .36 | .65 | large |
|  | DN |  | 94 | 3368 | 5.93e-06 | 1.78e-05 |  | .47 | .29 | .64 | moderate |
| IN |  | 99 |  | 4171 | 2.83e-14 | 8.49e-14 |  | .79 | .70 | .86 | large |

**Table** **S5**. Nonparametric pairwise comparisons using Wilcoxon signed-rank test investigating interactions between **(A)** direction and lobule; **(B)** direction and segment; and **(C)** lobule and segment. All confidence intervals are bootstrapped using 10,000 iterations to yield stable estimates around the effect size.

| **Polysynaptic Tractography Interactions: Controlling for Average Streamline Length** | | | | | | | | | | | | |
| --- | --- | --- | --- | --- | --- | --- | --- | --- | --- | --- | --- | --- |
| **A. Direction x Lobule** | | | | | | | | | | | | |
| Crus I | | | | | | | | | | | | |
| Measure 1 | Measure 2 | *n* 1 | *n* 2 | W | | *p* | *p_bonferroni_* | | *Eff. Size* | CI 95% | | Magnitude |
|  |  |  |  |  |  |  |  |  |  | Lower | Upper |  |
| R-R | L-L | 97 | 87 | 3125 | | 2.32e-09 | 1.39e-08 | **** | .65 | .51 | .77 | large |
|  | R-L |  | 90 | 3962 | | 1.10e-15 | 6.60e-15 |  | .85 | .82 | .87 |  |
|  | L-R |  | 52 | 1326 | | 5.30e-10 | 3.18e-09 |  | .87 | .87 | .87 |  |
| L-L | R-L | 87 | 90 | 2835 | | 3.25e-08 | 1.95e-07 |  | .61 | .46 | .74 |  |
|  | L-R |  | 52 | 1325 | | 5.63e-10 | 3.38e-09 |  | .87 | .86 | .87 |  |
| R-L |  | 90 | 52 | 1237 | | 7.63e-08 | 4.58e-07 |  | .75 | .61 | .85 |  |
| Crus II | | | | | | | | | | | | |
| Measure 1 | Measure 2 | *n* 1 | *n* 2 | W | | *p* | *p_bonferroni_* | | *Eff. Size* | CI 95% | | Magnitude |
|  |  |  |  |  |  |  |  |  |  | Lower | Upper |  |
| R-R | L-L | 99 | 90 | 3195 | | 3.93e-06 | 2.36e-05 | **** | .49 | 0.32 | 0.64 | moderate |
|  | R-L |  | 88 | 3864 | | 2.22e-15 | 1.33e-14 |  | .85 | 0.81 | 0.86 | large |
|  | L-R |  | 60 | 1830 | | 1.67e-11 | 1.00e-10 |  | .87 | 0.87 | 0.87 |  |
| L-L | R-L | 90 | 88 | 3029 | | 8.54e-10 | 5.12e-09 |  | .68 | 0.54 | 0.78 |  |
|  | L-R |  | 60 | 1828 | | 1.85e-11 | 1.11e-10 |  | .87 | 0.86 | 0.87 |  |
| R-L |  | 88 |  | 1460 | | 6.82e-08 | 4.09e-07 |  | .72 | 0.56 | 0.84 |  |
| Lobule VI | | | | | | | | | | | | |
| Measure 1 | Measure 2 | *n* 1 | *n* 2 | W | | *p* | *p_bonferroni_* | | *Eff. Size* | CI 95% | | Magnitude |
|  |  |  |  |  |  |  |  |  |  | Lower | Upper |  |
| R-R | L-L | 95 | 86 | 2573 | | 4.45e-04 | 3.00e-03 | ** | .38 | .19 | .57 | moderate |
|  | R-L |  | 88 | 3888 | | 9.88e-16 | 5.93e-15 | **** | .86 | .83 | .87 | large |
|  | L-R |  | 45 | 990 | | 1.14e-13 | 6.84e-13 |  | .87 | .87 | .87 |  |
| L-L | R-L | 86 | 88 | 3120 | | 6.45e-12 | 3.87e-11 |  | .76 | .67 | .83 |  |
|  | L-R |  | 45 | 989 | | 2.27e-13 | 1.36e-12 |  | .87 | .86 | .87 |  |
| R-L |  | 88 |  | 808 | | 1.87e-05 | 1.12e-04 | *** | .62 | .41 | .78 |  |
| **B. Direction x Segment** | | | | | | | | | | | | |
| a | | | | | | | | | | | | |
| Measure 1 | Measure 2 | *n* 1 | *n* 2 | W | | *p* | *p_bonferroni_* | | *Eff. Size* | CI 95% | | Magnitude |
|  |  |  |  |  |  |  |  |  |  | Lower | Upper |  |
| R-R | L-L | 100 | 97 | 3694 | | 6.04e-07 | 3.62e-06 | **** | .51 | .34 | .67 | large |
|  | R-L |  | 98 | 4659 | | 2.19e-16 | 1.31e-15 |  | .83 | .79 | .86 |  |
|  | L-R |  | 78 | 3002 | | 2.61e-14 | 1.57e-13 |  | .87 | .87 | .87 |  |
| L-L | R-L | 97 | 98 | 3517 | | 1.40e-05 | 8.40e-05 |  | .44 | .26 | .60 | moderate |
|  | L-R |  | 78 | 3080 | | 1.78e-14 | 1.07e-13 |  | .87 | .87 | .87 | large |
| R-L |  | 98 |  | 2706 | | 6.53e-09 | 3.92e-08 |  | .66 | .51 | .78 |  |
| b | | | | | | | | | | | | |
| Measure 1 | Measure 2 | *n* 1 | *n* 2 | W | | *p* | *p_bonferroni_* | | *Eff. Size* | CI 95% | | Magnitude |
|  |  |  |  |  |  |  |  |  |  | Lower | Upper |  |
| R-R | L-L | 101 | 100 | 3577 | | 3.00e-04 | 2.00e-03 | ** | .36 | .18 | .53 | moderate |
|  | R-L |  | 98 | 4773 | | 9.04e-17 | 5.42e-16 | **** | .84 | .81 | .86 | large |
|  | L-R |  | 85 | 3655 | | 1.19e-15 | 7.14e-15 |  | .87 | .87 | .87 |  |
| L-L | R-L | 100 | 98 | 4244 | | 1.84e-11 | 1.10e-10 |  | .68 | .57 | .78 |  |
|  | L-R |  | 85 | 3653 | | 1.28e-15 | 7.68e-15 |  | .87 | .86 | .87 |  |
| R-L |  | 98 |  | 3144 | | 2.04e-10 | 1.22e-09 |  | .70 | .57 | .80 |  |
| c_1_ | | | | | | | | | | | | |
| Measure 1 | Measure 2 | *n* 1 | *n* 2 | W | | *p* | *p_bonferroni_* | | *Eff. Size* | CI 95% | | Magnitude |
|  |  |  |  |  |  |  |  |  |  | Lower | Upper |  |
| R-R | L-L | 99 | 94 | 3302 | | 1.90e-05 | 1.14e-04 | *** | .44 | .26 | .61 | moderate |
|  | R-L |  | 96 | 4627 | | 4.49e-17 | 2.69e-16 | **** | .86 | .84 | .87 | large |
|  | L-R |  | 76 | 2926 | | 3.68e-14 | 2.21e-13 |  | .87 | .87 | .87 |  |
| L-L | R-L | 94 | 96 | 3441 | | 4.02e-07 | 2.41e-06 |  | .53 | .36 | .66 |  |
|  | L-R |  | 76 | 2925 | | 3.83e-14 | 2.30e-13 |  | .87 | .87 | .87 |  |
| R-L |  | 96 |  | 2643 | | 1.28e-10 | 7.68e-10 |  | .74 | .63 | .83 |  |
| c_2_ | | | | | | | | | | | | |
| Measure 1 | Measure 2 | *n* 1 | *n* 2 | | W | *p* | *p_bonferroni_* | | *Eff. Size* | CI 95% | | Magnitude |
|  |  |  |  |  |  |  |  |  |  | Lower | Upper |  |
| R-R | L-L | 99 | 91 | 3404 | | 4.87e-08 | 2.92e-07 | **** | .58 | .41 | .72 | large |
|  | R-L |  | 92 | 4250 | | 2.07e-16 | 1.24e-15 |  | .86 | .84 | .87 |  |
|  | L-R |  | 64 | 2079 | | 3.79e-12 | 2.27e-11 |  | .87 | .86 | .87 |  |
| L-L | R-L | 91 | 92 | 2962 | | 9.27e-06 | 5.56e-05 |  | .48 | .30 | .63 | moderate |
|  | L-R |  | 64 | 2077 | | 4.16e-12 | 2.50e-11 |  | .87 | .86 | .87 | large |
| R-L |  | 92 |  | 1825 | | 2.76e-09 | 1.66e-08 |  | .76 | .62 | .85 |  |
| c_3_ | | | | | | | | | | | | |
| Measure 1 | Measure 2 | *n* 1 | *n* 2 | W | | *p* | *p_bonferroni_* | | *Eff. Size* | CI 95% | | Magnitude |
|  |  |  |  |  |  |  |  |  |  | Lower | Upper |  |
| R-R | L-L | 98 | 87 | 3065 | | 2.73e-07 | 1.64e-06 | **** | .55 | .38 | .71 | large |
|  | R-L |  | 91 | 4064 | | 4.99e-16 | 2.99e-15 |  | .86 | .83 | .87 |  |
|  | L-R |  | 50 | 1224 | | 7.11e-15 | 4.27e-14 |  | .87 | .86 | .87 |  |
| L-L | R-L | 87 | 91 | 2679 | | 6.28e-06 | 3.77e-05 |  | .50 | .32 | .66 | moderate |
|  | L-R |  | 50 | 1222 | | 1.78e-14 | 1.07e-13 |  | .87 | .85 | .87 | large |
| R-L |  | 91 |  | 1101 | | 1.23e-07 | 7.38e-07 |  | .69 | .51 | .83 |  |
| c_4_ | | | | | | | | | | | | |
| Measure 1 | Measure 2 | *n* 1 | *n* 2 | W | | *p* | *p_bonferroni_* | | *Eff. Size* | CI 95% | | Magnitude |
|  |  |  |  |  |  |  |  |  |  | Lower | Upper |  |
| R-R | L-L | 97 | 88 | 2639 | | 9.43e-04 | 6.00e-03 | ** | .36 | .16 | .54 | moderate |
|  | R-L |  |  | 3879 | | 1.34e-15 | 8.04e-15 | **** | .85 | .82 | .87 | large |
|  | L-R |  | 59 | 1761 | | 3.89e-11 | 2.33e-10 |  | .86 | .84 | .87 |  |
| L-L | R-L | 88 | 88 | 2533 | | 1.21e-05 | 7.26e-05 |  | .49 | .30 | .65 | moderate |
|  | L-R |  | 59 | 1767 | | 2.86e-11 | 1.72e-10 |  | .87 | .86 | .87 | large |
| R-L |  |  |  | 1430 | | 2.59e-07 | 1.55e-06 |  | .69 | .52 | .82 |  |
| **C. Lobule x Segment** | | | | | | | | | | | | |
| a | | | | | | | | | | | | |
| Measure 1 | Measure 2 | *n* 1 | *n* 2 | W | | *p* | *p_bonferroni_* | | *Eff. Size* | CI 95% | | Magnitude |
|  |  |  |  |  |  |  |  |  |  | Lower | Upper |  |
| Crus I | Crus II | 80 | 90 | 440 | | 7.16e-08 | 2.15e-07 | **** | .61 | .45 | .76 | large |
|  | Lobule VI |  | 98 | 22 | | 1.83e-14 | 5.49e-14 |  | .86 | .83 | .87 |  |
| Crus II |  | 90 |  | 317 | | 3.38e-12 | 1.01e-11 |  | .73 | .63 | .81 |  |
| b | | | | | | | | | | | | |
| Measure 1 | Measure 2 | *n* 1 | *n* 2 | W | | *p* | *p_bonferroni_* | | *Eff. Size* | CI 95% | | Magnitude |
|  |  |  |  |  |  |  |  |  |  | Lower | Upper |  |
| Crus I | Crus II | 91 | 94 | 3078 | | 3.19e-06 | 9.57e-06 | **** | .50 | .32 | .64 | moderate |
|  | Lobule VI |  | 89 | 2109 | | 2.18e-01 | 6.54e-01 | ns | .13 | .007 | .34 | small |
| Crus II |  | 94 |  | 1319 | | 1.80e-02 | 5.30e-02 |  | .26 | .05 | .45 |  |
| c_1_ | | | | | | | | | | | | |
| Measure 1 | Measure 2 | *n* 1 | *n* 2 | W | | *p* | *p_bonferroni_* | | *Eff. Size* | CI 95% | | Magnitude |
|  |  |  |  |  |  |  |  |  |  | Lower | Upper |  |
| Crus I | Crus II | 94 | 89 | 2480 | | .009 | .026 | * | .28 | .08 | .47 | small |
|  | Lobule VI |  | 77 | 2069 | | .002 | .005 | ** | .36 | .14 | .57 | moderate |
| Crus II |  | 89 |  | 1549 | | .514 | > .999 | ns | .08 | .004 | .31 | small |
| c_2_ | | | | | | | | | | | | |
| Measure 1 | Measure 2 | *n* 1 | *n* 2 | W | | *p* | *p_bonferroni_* | | *Eff. Size* | CI 95% | | Magnitude |
|  |  |  |  |  |  |  |  |  |  | Lower | Upper |  |
| Crus I | Crus II | 83 | 87 | 963 | | 3.00e-03 | 8.00e-03 | ** | .34 | .13 | .53 | moderate |
|  | Lobule VI |  | 64 | 2016 | | 5.29e-12 | 1.59e-11 | **** | .87 | .87 | .87 | large |
| Crus II |  | 87 |  | 1944 | | 1.20e-11 | 3.60e-11 |  | .86 | .84 | .87 |  |
| c_3_ | | | | | | | | | | | | |
| Measure 1 | Measure 2 | *n* 1 | *n* 2 | W | | *p* | *p_bonferroni_* | | *Eff. Size* | CI 95% | | Magnitude |
|  |  |  |  |  |  |  |  |  |  | Lower | Upper |  |
| Crus I | Crus II | 77 | 77 | 467 | | 5.75e-06 | 1.72e-05 | **** | .54 | .36 | .70 | large |
|  | Lobule VI |  | 50 | 1176 | | 7.11e-15 | 2.13e-14 |  | .87 | .87 | .87 |  |
| Crus II |  |  |  | 1275 | | 7.79e-10 | 2.34e-09 |  |  |  |  |  |
| c_4_ | | | | | | | | | | | | |
| Measure 1 | Measure 2 | *n* 1 | *n* 2 | W | | *p* | *p_bonferroni_* | | *Eff. Size* | CI 95% | | Magnitude |
|  |  |  |  |  |  |  |  |  |  | Lower | Upper |  |
| Crus I | Crus II | 64 | 69 | 679 | | 0.334 | - | ns | .13 | .007 | .38 | small |

The following are the results from the main analysis repeated when controlling for volume of the DCN—another potential confound that may affect streamline results for the polysynaptic circuit.

**Table S6.** Nonparametric pairwise comparisons using Wilcoxon sign-rank test investigating streamline count differences based on tract direction, lobule, and segment for polysynaptic tractographies.

| **Polysynaptic Tractography: Controlling for Waypoint/DCN Volume** | | | | | | | | | | | |
| --- | --- | --- | --- | --- | --- | --- | --- | --- | --- | --- | --- |
| Direction | | | | | | | | | | | |
| Measure 1 | Measure 2 | *n* 1 | *n* 2 | W | *p* | *p_bonferroni_* | | *Eff. Size* | CI 95% | | Magnitude |
|  |  |  |  |  |  |  |  |  | Lower | Upper |  |
| R-R | L-L | 92 | 80 | 2363 | 3.21e-06 | 1.93e-05 | **** | .53 | .35 | .70 | large |
|  | R-L |  |  | 3123 | 4.76e-14 | 2.86e-13 |  | .85 | .82 | .87 |  |
|  | L-R |  | 32 | 496 | 9.31e-10 | 5.59e-09 |  | .87 | .87 | .87 |  |
| L-L | R-L | 80 | 80 | 2208 | 1.00e-07 | 6.00e-07 |  | .63 | .47 | .76 |  |
|  | L-R |  | 32 | 527 | 9.31e-10 | 5.59e-09 |  | .87 | .85 | .87 |  |
| R-L |  |  |  | 444 | 3.57e-05 | 2.14e-04 | *** | .69 | .45 | .85 |  |
| Lobule | | | | | | | | | | | |
| Measure 1 | Measure 2 | *n* 1 | *n* 2 | W | *p* | *p_bonferroni_* | | *Eff. Size* | CI 95% | | Magnitude |
|  |  |  |  |  |  |  |  |  | Lower | Upper |  |
| Crus I | Crus II | 49 | 56 | 472 | 0.599 | > .999 | ns | .08 | .005 | .38 | small |
|  | Lobule VI |  | 43 | 139 | 0.003 | 0.01 | ** | .49 | .19 | .73 | moderate |
| Crus II |  | 56 |  | 190 | 0.024 | 0.072 | † | .37 | .07 | .64 |  |
| Segment | | | | | | | | | | | |
| Measure 1 | Measure 2 | *n* 1 | *n* 2 | W | *p* | *p_bonferroni_* | | *Eff. Size* | CI 95% | | Magnitude |
|  |  |  |  |  |  |  |  |  | Lower | Upper |  |
| a | b | 77 | 83 | 2237 | 3.97e-08 | 5.96e-07 | **** | .65 | .49 | .78 | large |
|  | c_1_ |  | 75 | 1934 | 1.95e-11 | 2.92e-10 |  | .85 | .81 | .87 |  |
|  | c_2_ |  | 62 | 1223 | 1.07e-14 | 1.60e-13 |  | .87 | .85 |  |  |
|  | c_3_ |  | 48 | 703 | 1.46e-11 | 2.19e-10 |  | .87 | .87 |  |  |
|  | c_4_ |  | 56 | 946 | 2.27e-13 | 3.40e-12 |  | .87 | .87 |  |  |
| b | c_1_ | 83 | 75 | 2673 | 3.65e-13 | 5.48e-12 |  | .85 | .81 |  |  |
|  | c_2_ |  | 62 | 1758 | 4.53e-11 | 6.80e-10 |  | .86 | .83 |  |  |
|  | c_3_ |  | 48 | 1128 | 1.42e-14 | 2.13e-13 |  | .87 | .87 |  |  |
|  | c_4_ |  | 56 | 1431 | 2.46e-10 | 3.69e-09 |  | .87 | .87 |  |  |
| c_1_ | c_2_ | 75 | 62 | 1667 | 3.66e-09 | 5.49e-08 |  | .77 | .66 | 0.84 |  |
|  | c_3_ |  | 48 | 1126 | 4.26e-14 | 6.39e-13 |  | .87 | .85 | .87 |  |
|  | c_4_ |  | 56 | 1485 | 1.67e-10 | 2.50e-09 |  |  | .87 |  |  |
| c_2_ | c_3_ | 62 | 48 | 990 | 1.14e-13 | 1.71e-12 |  |  |  |  |  |
|  | c_4_ |  | 56 | 1225 | 3.55e-15 | 5.32e-14 |  |  |  |  |  |
| c_3_ |  | 48 |  | 946 | 2.27e-13 | 3.40e-12 |  |  |  |  |  |

Table **S7**. Nonparametric pairwise comparisons using Wilcoxon signed-rank test investigating interactions between **(A)** direction and lobule; **(B)** direction and segment; and **(C)** lobule and segment. All confidence intervals are bootstrapped using 10,000 iterations to yield stable estimates around the effect size.

| Polysynaptic Tractography Interactions: Controlling for Waypoint/DCN Volume | | | | | | | | | | | | |
| --- | --- | --- | --- | --- | --- | --- | --- | --- | --- | --- | --- | --- |
| **A. Direction x Lobule** | | | | | | | | | | | | |
| Crus I | | | | | | | | | | | | |
| Measure 1 | Measure 2 | *n* 1 | *n* 2 | W | | *p* | *p_bonferroni_* | | *Eff. Size* | CI 95% | | Magnitude |
|  |  |  |  |  |  |  |  |  |  | Lower | Upper |  |
| R-R | L-L | 97 | 87 | 3099 | | 4.69e-09 | 2.81e-08 | **** | .64 | .49 | .76 | large |
|  | R-L |  | 90 | 3938 | | 2.44e-15 | 1.46e-14 |  | .84 | .80 | .86 |  |
|  | L-R |  | 52 | 1326 | | 5.30e-10 | 3.18e-09 |  | .87 | .87 | .87 |  |
| L-L | R-L | 87 | 90 | 2707 | | 8.44e-07 | 5.06e-06 |  | .55 | .37 | .70 |  |
|  | L-R |  | 52 | 1325 | | 5.63e-10 | 3.38e-09 |  | .87 | .86 | .87 |  |
| R-L |  | 90 | 52 | 1235 | | 8.46e-08 | 5.08e-07 |  | .75 | .61 | .85 |  |
| Crus II | | | | | | | | | | | | |
| Measure 1 | Measure 2 | *n* 1 | *n* 2 | W | | *p* | *p_bonferroni_* | | *Eff. Size* | CI 95% | | Magnitude |
|  |  |  |  |  |  |  |  |  |  | Lower | Upper |  |
| R-R | L-L | 99 | 90 | 3223 | | 2.27e-06 | 1.36e-05 | **** | .50 | .33 | .65 | moderate |
|  | R-L |  | 88 | 3838 | | 5.27e-15 | 3.16e-14 |  | .83 | .79 | .86 | large |
|  | L-R |  | 60 | 1829 | | 1.76e-11 | 1.06e-10 |  | .87 | .86 | .87 |  |
| L-L | R-L | 90 | 88 | 2907 | | 2.54e-08 | 1.52e-07 |  | .62 | .46 | .75 |  |
|  | L-R |  | 60 | 1828 | | 1.85e-11 | 1.11e-10 |  | .87 | .86 | .87 |  |
| R-L |  | 88 |  | 1452 | | 9.79e-08 | 5.87e-07 |  | .71 | .56 | .83 |  |
| Lobule VI | | | | | | | | | | | | |
| Measure 1 | Measure 2 | *n* 1 | *n* 2 | W | | *p* | *p_bonferroni_* | | *Eff. Size* | CI 95% | | Magnitude |
|  |  |  |  |  |  |  |  |  |  | Lower | Upper |  |
| R-R | L-L | 95 | 86 | 2663 | | 9.10e-05 | 5.46e-04 | *** | .43 | .24 | .61 | moderate |
|  | R-L |  | 88 | 3861 | | 2.45e-15 | 1.47e-14 | ****  *** | .84 | .81 | .86 | large |
|  | L-R |  | 45 | 990 | | 1.14e-13 | 6.84e-13 |  | .87 | .87 | .87 |  |
| L-L | R-L | 86 | 88 | 3043 | | 7.67e-11 | 4.60e-10 |  | .72 | .61 | .81 |  |
|  | L-R |  | 45 | 989 | | 2.27e-13 | 1.36e-12 |  | .87 | .86 | .87 |  |
| R-L |  | 88 |  | 801 | | 2.97e-05 | 1.78e-04 |  | .60 | .38 | .78 |  |
| B. Direction x Segment | | | | | | | | | | | | |
| a | | | | | | | | | | | | |
| Measure 1 | Measure 2 | *n* 1 | *n* 2 | W | | *p* | *p_bonferroni_* | | *Eff. Size* | CI 95% | | Magnitude |
|  |  |  |  |  |  |  |  |  |  | Lower | Upper |  |
| R-R | L-L | 100 | 97 | 3711 | | 4.37e-07 | 2.62e-06 | **** | .52 | .35 | .67 | large |
|  | R-L |  | 98 | 4644 | | 3.43e-16 | 2.06e-15 |  | .83 | .78 | .86 |  |
|  | L-R |  | 78 | 3002 | | 2.61e-14 | 1.57e-13 |  | .87 | .87 | .87 |  |
| L-L | R-L | 97 | 98 | 3452 | | 4.03e-05 | 2.42e-04 | *** | .42 | .24 | .58 | moderate |
|  | L-R |  | 78 | 3080 | | 1.78e-14 | 1.07e-13 | **** | .87 | .87 | .87 | large |
| R-L |  | 98 |  | 2738 | | 2.49e-09 | 1.49e-08 |  | .68 | .53 | .79 |  |
| b | | | | | | | | | | | | |
| Measure 1 | Measure 2 | *n* 1 | *n* 2 | W | | *p* | *p_bonferroni_* | | *Eff. Size* | CI 95% | | Magnitude |
|  |  |  |  |  |  |  |  |  |  | Lower | Upper |  |
| R-R | L-L | 101 | 100 | 3580 | | 2.88e-04 | 2.00e-03 | ** | .36 | .18 | .53 | moderate |
|  | R-L |  | 98 | 4764 | | 1.18e-16 | 7.08e-16 | **** | .84 | .80 | .86 | large |
|  | L-R |  | 85 | 3655 | | 1.19e-15 | 7.14e-15 |  | .87 | .87 | .87 |  |
| L-L | R-L | 100 | 98 | 4224 | | 3.01e-11 | 1.81e-10 |  | .67 | .56 | .77 |  |
|  | L-R |  | 85 | 3653 | | 1.28e-15 | 7.68e-15 |  | .87 | .86 | .87 |  |
| R-L |  | 98 |  | 3159 | | 1.31e-10 | 7.86e-10 |  | .71 | .58 | .80 |  |
| c_1_ | | | | | | | | | | | | |
| Measure 1 | Measure 2 | *n* 1 | *n* 2 | W | | *p* | *p_bonferroni_* | | *Eff. Size* | CI 95% | | Magnitude |
|  |  |  |  |  |  |  |  |  |  | Lower | Upper |  |
| R-R | L-L | 99 | 94 | 3360 | | 6.85e-06 | 4.11e-05 | **** | .47 | .29 | .63 | moderate |
|  | R-L |  | 96 | 4618 | | 5.93e-17 | 3.56e-16 |  | .85 | .83 | .87 | large |
|  | L-R |  | 76 | 2926 | | 3.68e-14 | 2.21e-13 |  | .87 | .87 | .87 |  |
| L-L | R-L | 94 | 96 | 3438 | | 4.28e-07 | 2.57e-06 |  | .53 | .36 | .66 |  |
|  | L-R |  | 76 | 2925 | | 3.83e-14 | 2.30e-13 |  | .87 | .87 | .87 |  |
| R-L |  | 96 |  | 2669 | | 5.16e-11 | 3.10e-10 |  | .76 | .65 | .84 |  |
| c_2_ | | | | | | | | | | | | |
| Measure 1 | Measure 2 | *n* 1 | *n* 2 | | W | *p* | *p_bonferroni_* | | *Eff. Size* | CI 95% | | Magnitude |
|  |  |  |  |  |  |  |  |  |  | Lower | Upper |  |
| R-R | L-L | 99 | 91 | 3434 | | 2.45e-08 | 1.47e-07 | **** | .59 | .43 | .73 | large |
|  | R-L |  | 92 | 4238 | | 3.05e-16 | 1.83e-15 |  | .85 | .83 | .87 |  |
|  | L-R |  | 64 | 2079 | | 3.79e-12 | 2.27e-11 |  | .87 | .86 | .87 |  |
| L-L | R-L | 91 | 92 | 2962 | | 9.27e-06 | 5.56e-05 |  | .48 | .30 | .63 | moderate |
|  | L-R |  | 64 | 2077 | | 4.16e-12 | 2.50e-11 |  | .87 | .86 | .87 | large |
| R-L |  | 92 |  | 1829 | | 2.32e-09 | 1.39e-08 |  | .76 | .63 | .85 |  |
| c_3_ | | | | | | | | | | | | |
| Measure 1 | Measure 2 | *n* 1 | *n* 2 | W | | *p* | *p_bonferroni_* | | *Eff. Size* | CI 95% | | Magnitude |
|  |  |  |  |  |  |  |  |  |  | Lower | Upper |  |
| R-R | L-L | 98 | 87 | 3076 | | 2.12e-07 | 1.27e-06 | **** | .56 | .39 | .71 | large |
|  | R-L |  | 91 | 4061 | | 5.51e-16 | 3.31e-15 |  | .85 | .83 | .87 |  |
|  | L-R |  | 50 | 1224 | | 7.11e-15 | 4.27e-14 |  | .87 | .86 | .87 |  |
| L-L | R-L | 87 | 91 | 2667 | | 8.15e-06 | 4.89e-05 |  | .49 | .31 | .65 | moderate |
|  | L-R |  | 50 | 1222 | | 1.78e-14 | 1.07e-13 |  | .87 | .85 | .87 | large |
| R-L |  | 91 |  | 1107 | | 7.75e-08 | 4.65e-07 |  | .70 | .53 | .83 |  |
| c_4_ | | | | | | | | | | | | |
| Measure 1 | Measure 2 | *n* 1 | *n* 2 | W | | *p* | *p_bonferroni_* | | *Eff. Size* | CI 95% | | Magnitude |
|  |  |  |  |  |  |  |  |  |  | Lower | Upper |  |
| R-R | L-L | 97 | 88 | 2681 | | 4.87e-04 | 3.00e-03 | ** | .38 | .18 | .55 | moderate |
|  | R-L |  |  | 3796 | | 1.68e-15 | 1.01e-14 | **** | .85 | .82 | .87 | large |
|  | L-R |  | 59 | 1763 | | 3.51e-11 | 2.11e-10 |  | .86 | .84 | .87 |  |
| L-L | R-L | 88 | 88 | 2526 | | 1.41e-05 | 8.46e-05 |  | .49 | .31 | .64 | moderate |
|  | L-R |  | 59 | 1767 | | 2.86e-11 | 1.72e-10 |  | .87 | .86 | .87 | large |
| R-L |  |  |  | 1440 | | 1.67e-07 | 1.00e-06 |  | .70 | .54 | .82 |  |
| C. Lobule x Segment | | | | | | | | | | | | |
| a | | | | | | | | | | | | |
| Measure 1 | Measure 2 | *n* 1 | *n* 2 | W | | *p* | *p_bonferroni_* | | *Eff. Size* | CI 95% | | Magnitude |
|  |  |  |  |  |  |  |  |  |  | Lower | Upper |  |
| Crus I | Crus II | 80 | 90 | 530 | | 8.21e-07 | 2.46e-06 | **** | .56 | .38 | .72 | large |
|  | Lobule VI |  | 98 | 80 | | 1.54e-13 | 4.62e-13 |  | .83 | .77 | .86 |  |
| Crus II |  | 90 |  | 472 | | 2.34e-10 | 7.02e-10 |  | .67 | .53 | .78 |  |
| b | | | | | | | | | | | | |
| Measure 1 | Measure 2 | *n* 1 | *n* 2 | W | | *p* | *p_bonferroni_* | | *Eff. Size* | CI 95% | | Magnitude |
|  |  |  |  |  |  |  |  |  |  | Lower | Upper |  |
| Crus I | Crus II | 91 | 94 | 3000 | | 1.47e-05 | 4.41e-05 | **** | .46 | .28 | .62 | moderate |
|  | Lobule VI |  | 89 | 2724 | | 8.63e-05 | 2.59e-04 | *** | .43 | .23 | .60 |  |
| Crus II |  | 94 |  | 1725 | | 5.32e-01 | > .999 | ns | .07 | .004 | .28 | small |
| c_1_ | | | | | | | | | | | | |
| Measure 1 | Measure 2 | *n* 1 | *n* 2 | W | | *p* | *p_bonferroni_* | | *Eff. Size* | CI 95% | | Magnitude |
|  |  |  |  |  |  |  |  |  |  | Lower | Upper |  |
| Crus I | Crus II | 94 | 89 | 2314 | | 5.60e-02 | 1.69e-01 | ns | .21 | .02 | .40 | small |
|  | Lobule VI |  | 77 | 2473 | | 1.73e-07 | 5.19e-07 | **** | .60 | .43 | .75 | large |
| Crus II |  | 89 |  | 1954 | | 5.00e-03 | 1.60e-02 | * | .32 | .11 | .53 | moderate |
| c_2_ | | | | | | | | | | | | |
| Measure 1 | Measure 2 | *n* 1 | *n* 2 | W | | *p* | *p_bonferroni_* | | *Eff. Size* | CI 95% | | Magnitude |
|  |  |  |  |  |  |  |  |  |  | Lower | Upper |  |
| Crus I | Crus II | 83 | 87 | 986 | | 4.00e-03 | 1.10e-02 | * | .33 | .12 | .52 | moderate |
|  | Lobule VI |  | 64 | 2016 | | 5.29e-12 | 1.59e-11 | **** | .87 | .87 | .87 | large |
| Crus II |  | 87 |  | 1952 | | 8.16e-12 | 2.45e-11 |  | .87 | .86 | .87 |  |
| c_3_ | | | | | | | | | | | | |
| Measure 1 | Measure 2 | *n* 1 | *n* 2 | W | | *p* | *p_bonferroni_* | | *Eff. Size* | CI 95% | | Magnitude |
|  |  |  |  |  |  |  |  |  |  | Lower | Upper |  |
| Crus I | Crus II | 77 | 77 | 538 | | 3.79e-05 | 1.14e-04 | *** | .49 | .29 | .66 | moderate |
|  | Lobule VI |  | 50 | 1176 | | 7.11e-15 | 2.13e-14 | **** | .87 | .87 | .87 | large |
| Crus II |  |  |  | 1275 | | 7.79e-10 | 2.34e-09 |  |  |  |  |  |
| c_4_ | | | | | | | | | | | | |
| Measure 1 | Measure 2 | *n* 1 | *n* 2 | W | | *p* | *p_bonferroni_* | | *Eff. Size* | CI 95% | | Magnitude |
|  |  |  |  |  |  |  |  |  |  | Lower | Upper |  |
| Crus I | Crus II | 64 | 69 | 713 | | .491 | - | ns | .09 | .006 | .35 | small |

**Table S8.** **(A)** Average and segment-wise correlations with the NEO Five Factor Inventory (NEO-FFI) subscales for Neuroticism and Agreeableness and fractional anisotropy (FA) values for the right-to-right travelling tracts with the highest densities. The N for all correlations is 101, and all confidence intervals are bootstrapped using 10,000 iterations. False Discovery Rate (FDR) correction for multiple comparisons was performed for the number of tracts within the polysynaptic (4 tracts) and monosynaptic (3 tracts) correlation domains for overall tract averages, separately. Corrections were not performed for the 4 quarters, as this was an exploratory analysis used to “drill down” into which portions of the tract were driving the effects where significant or trending associations were uncovered. Note that there was a significant difference between the sexes for each personality trait, so partial correlations are reported in section **(B)**.

| **A. Fractional Anisotropy Correlations with Personality Factors** | | | | | | | | | | |
| --- | --- | --- | --- | --- | --- | --- | --- | --- | --- | --- |
| **Big Five Factor Inventory Neuroticism** | | | | | | | | | | |
| **Type** | **Lobule** | **Segment/DCN** | **Section** | **Rho (*ρ*)** | ***p*-value** | | ***p*-value FDR** | | **95% CI** | |
|  |  |  |  |  |  |  |  |  | **Lower** | **Upper** |
| R-R Poly-synaptic | VI | a (vermis) | **Avg.** | **.28** | **.005** | ****** | **.012** | ***** | **.07** | **.46** |
|  |  |  | *Q-1* | .13 | .212 | ns |  | | -.08 | .32 |
|  |  |  | *Q-2* | .21 | .032 | * |  |  | .01 | .40 |
|  |  |  | *Q-3* | .25 | .010 | * |  |  | .05 | .44 |
|  |  |  | *Q-4* | .14 | .171 | ns |  |  | -.07 | .33 |
|  | Crus I | b (paravermis) | **Avg.** | **.27** | **.006** | ****** | **.012** | ***** | **.08** | **.45** |
|  |  |  | *Q-1* | .14 | .172 | ns |  | | -.06 | .33 |
|  |  |  | *Q-2* | .29 | .004 | ** |  |  | .09 | .46 |
|  |  |  | *Q-3* | .24 | .015 | * |  |  | .04 | .42 |
|  |  |  | *Q-4* | .13 | .189 | ns |  |  | -.08 | .33 |
|  |  | c_1_ (cerebro-cerebellum) | **Avg.** | **.22** | **.026** | ***** | **.026** | ***** | **.02** | **.40** |
|  |  |  | *Q-1* | .04 | .712 | ns |  | | -.16 | .25 |
|  |  |  | *Q-2* | .14 | .169 | ns |  |  | -.05 | .32 |
|  |  |  | *Q-3* | .21 | .039 | * |  |  | .01 | .39 |
|  |  |  | *Q-4* | .13 | .196 | ns |  |  | -.08 | .33 |
|  | Crus II |  | **Avg.** | **.25** | **.012** | ***** | **.016** | ***** | **.05** | **.43** |
|  |  |  | *Q-1* | .19 | .059 | † |  | | -.004 | .37 |
|  |  |  | *Q-2* | .15 | .142 | ns |  |  | -.05 | .34 |
|  |  |  | *Q-3* | .21 | .040 | * |  |  | .01 | .39 |
|  |  |  | *Q-4* | .11 | .256 | ns |  |  | -.09 | .31 |
| R-R Mono-synaptic | Fastigial | | **Avg.** | **.25** | **.014** | ***** | **.036** | ***** | **.05** | **.42** |
|  |  |  | *Q-1* | .15 | .141 | ns |  | | -.06 | .34 |
|  |  |  | *Q-2* | .25 | .010 | * |  |  | .05 | .44 |
|  |  |  | *Q-3* | .18 | .065 | † |  |  | -.02 | .37 |
|  |  |  | *Q-4* | .08 | .432 | ns |  |  | -.13 | .28 |
|  | Interposed | | **Avg.** | **.22** | **.024** | ***** | **.036** | ***** | **.03** | **.40** |
|  |  |  | *Q-1* | .22 | .031 | * |  | | .02 | .39 |
|  |  |  | *Q-2* | .24 | .016 | * |  |  | .05 | .41 |
|  |  |  | *Q-3* | .13 | .186 | ns |  |  | -.07 | .32 |
|  |  |  | *Q-4* | .12 | .242 | ns |  |  | -.10 | .32 |
|  | Dentate | | **Avg.** | **.18** | **.074** | **†** | **.074** | **†** | **-.02** | **.37** |
|  |  |  | *Q-1* | .05 | .615 | ns |  | | -.15 | .24 |
|  |  |  | *Q-2* | .13 | .212 | ns |  |  | -.07 | .32 |
|  |  |  | *Q-3* | .15 | .129 | ns |  |  | -.04 | .34 |
|  |  |  | *Q-4* | .15 | .143 | ns |  |  | -.07 | .35 |
| **Big Five Factor Inventory Agreeableness** | | | | | | | | | | |
| **Type** | **Lobule** | **Segment/DCN** | **Section** | **Rho (*ρ*)** | ***p*-value** | | ***p*-value FDR** | | **95% CI** | |
|  |  |  |  |  |  |  |  |  | **Lower** | **Upper** |
| R-R Poly-synaptic | VI | a (vermis) | **Avg.** | **.24** | **.017** | ***** | **.068** | **†** | **.05** | **.41** |
|  |  |  | *Q-1* | .02 | .831 | ns |  | | -.18 | .23 |
|  |  |  | *Q-2* | .16 | .118 | ns |  |  | -.04 | .35 |
|  |  |  | *Q-3* | .29 | .003 | ** |  |  | .11 | .46 |
|  |  |  | *Q-4* | .17 | .084 | † |  |  | -.02 | .36 |
|  | Crus I | b (paravermis) | **Avg.** | **.17** | **.100** | **ns** | **.100** | **ns** | **-.03** | **.34** |
|  |  |  | *Q-1* | -.10 | .341 | ns |  | | -.30 | .11 |
|  |  |  | *Q-2* | -.02 | .838 | ns |  |  | -.22 | .19 |
|  |  |  | *Q-3* | .24 | .015 | * |  |  | .06 | .41 |
|  |  |  | *Q-4* | .14 | .156 | ns |  |  | -.05 | .33 |
|  |  | c_1_ (cerebro-cerebellum) | **Avg.** | **.17** | **.089** | **†** | **.100** | **ns** | **-.02** | **.35** |
|  |  |  | *Q-1* | -.16 | .120 | ns |  | | -.35 | .04 |
|  |  |  | *Q-2* | .04 | .707 | ns |  |  | -.17 | .24 |
|  |  |  | *Q-3* | .25 | .013 | * |  |  | .06 | .42 |
|  |  |  | *Q-4* | .13 | .185 | ns |  |  | -.06 | .32 |
|  | Crus II |  | **Avg.** | **.17** | **.096** | **†** | **.100** | **ns** | **-.01** | **.34** |
|  |  |  | *Q-1* | .01 | .951 | ns |  | | -.22 | .22 |
|  |  |  | *Q-2* | -.05 | .608 | ns |  |  | -.25 | .15 |
|  |  |  | *Q-3* | .20 | .041 | * |  |  | .02 | .38 |
|  |  |  | *Q-4* | .12 | .242 | ns |  |  | -.08 | .30 |
| R-R Mono-synaptic | Fastigial | | **Avg.** | **.27** | **.006** | ****** | **.015** | ***** | **.09** | **.44** |
|  |  |  | *Q-1* | .20 | .042 | * |  | | .01 | .38 |
|  |  |  | *Q-2* | .29 | .004 | ** |  |  | .10 | .45 |
|  |  |  | *Q-3* | .17 | .099 | † |  |  | -.03 | .35 |
|  |  |  | *Q-4* | .13 | .190 | ns |  |  | -.07 | .32 |
|  | Interposed | | **Avg.** | **.26** | **.010** | ****** | **.015** | ***** | **.08** | **.42** |
|  |  |  | *Q-1* | .22 | .025 | * |  | | .03 | .40 |
|  |  |  | *Q-2* | .26 | .008 | ** |  |  | .07 | .44 |
|  |  |  | *Q-3* | .15 | .134 | ns |  |  | -.04 | .33 |
|  |  |  | *Q-4* | .10 | .323 | ns |  |  | -.10 | .29 |
|  | Dentate | | **Avg.** | **.21** | **.033** | ***** | **.033** | ***** | **.02** | **.39** |
|  |  |  | *Q-1* | .04 | .728 | ns |  | | -.17 | .24 |
|  |  |  | *Q-2* | .15 | .124 | ns |  |  | -.05 | .35 |
|  |  |  | *Q-3* | .24 | .018 | * |  |  | .05 | .41 |
|  |  |  | *Q-4* | .09 | .377 | ns |  |  | -.11 | .28 |
| **B. Partial Correlations with Fractional Anisotropy, Personality Factors, and Gender** | | | | | | | | | | |
| **Big Five Factor Inventory Neuroticism** | | | | | | | | | | |
| **Type** | **Lobule** | **Segment/DCN** | **Section** | **Rho (*ρ*)** | ***p*-value** | | ***p*-value FDR** | | **95% CI** | |
|  |  |  |  |  |  |  |  |  | **Lower** | **Upper** |
| R-R Poly-synaptic | VI | a (vermis) | **Avg.** | **.24** | **.017** | ***** | **.041** | ***** | **.03** | **.43** |
|  |  |  | *Q-1* | .11 | .263 | ns |  | | -.09 | .31 |
|  |  |  | *Q-2* | .18 | .071 | † |  |  | -.02 | .38 |
|  |  |  | *Q-3* | .22 | .031 | * |  |  | .004 | .41 |
|  |  |  | *Q-4* | .11 | .262 | ns |  |  | -.10 | .32 |
|  | Crus I | b (paravermis) | **Avg.** | **.23** | **.024** | ***** | **.041** | ***** | **.02** | **.41** |
|  |  |  | *Q-1* | .12 | .223 | ns |  | | -.08 | .33 |
|  |  |  | *Q-2* | .24 | .016 | * |  |  | .06 | .42 |
|  |  |  | *Q-3* | .19 | .053 | † |  |  | -.01 | .38 |
|  |  |  | *Q-4* | .12 | .226 | ns |  |  | -.09 | .32 |
|  |  | c_1_ (cerebro-cerebellum) | **Avg.** | **.17** | **.087** | **†** | **.087** | **ns** | **-.03** | **.37** |
|  |  |  | *Q-1* | .02 | .841 | ns |  | | -.17 | .22 |
|  |  |  | *Q-2* | .08 | .435 | ns |  |  | -.11 | .27 |
|  |  |  | *Q-3* | .16 | .110 | ns |  |  | -.03 | .36 |
|  |  |  | *Q-4* | .12 | .239 | ns |  |  | -.09 | .32 |
|  | Crus II |  | **Avg.** | **.22** | **.031** | ***** | **.041** | ***** | **.03** | **.40** |
|  |  |  | *Q-1* | .19 | .064 | † |  | | -.01 | .38 |
|  |  |  | *Q-2* | .12 | .252 | ns |  |  | -.09 | .31 |
|  |  |  | *Q-3* | .17 | .095 | † |  |  | -.03 | .36 |
|  |  |  | *Q-4* | .10 | .325 | ns |  |  | -.11 | .30 |
| R-R Mono-synaptic | Fastigial | | **Avg.** | **.21** | **.037** | ***** | **.094** | **†** | **.01** | **.40** |
|  |  |  | *Q-1* | .11 | .262 | ns |  | | -.09 | .31 |
|  |  |  | *Q-2* | .21 | .032 | * |  |  | .02 | .41 |
|  |  |  | *Q-3* | .17 | .083 | † |  |  | -.03 | .37 |
|  |  |  | *Q-4* | .06 | .575 | ns |  |  | -.15 | .26 |
|  | Interposed | | **Avg.** | **.19** | **.063** | **†** | **.094** | **†** | ***.003*** | ***.37*** |
|  |  |  | *Q-1* | .16 | .108 | ns |  | | -.02 | .35 |
|  |  |  | *Q-2* | .20 | .051 | † |  |  | *.01* | *.38* |
|  |  |  | *Q-3* | .12 | .229 | ns |  |  | -.08 | .32 |
|  |  |  | *Q-4* | .10 | .333 | ns |  |  | -.11 | .31 |
|  | Dentate | | **Avg.** | **.13** | **.208** | **ns** | **.208** | **ns** | **-.07** | **.32** |
|  |  |  | *Q-1* | .002 | .986 | ns |  | | -.19 | .20 |
|  |  |  | *Q-2* | .07 | .472 | ns |  |  | -.13 | .27 |
|  |  |  | *Q-3* | .11 | .269 | ns |  |  | -.08 | .31 |
|  |  |  | *Q-4* | .14 | .163 | ns |  |  | -.07 | .35 |
| **Big Five Factor Inventory Agreeableness** | | | | | | | | | | |
| **Type** | **Lobule** | **Segment/DCN** | **Section** | **Rho (*ρ*)** | ***p*-value** | | ***p*-value FDR** | | **95% CI** | |
|  |  |  |  |  |  |  |  |  | **Lower** | **Upper** |
| R-R Poly-synaptic | VI | a (vermis) | **Avg.** | **.20** | **.042** | ***** | **.168** | **ns** | **.02** | **.38** |
|  |  |  | *Q-1* | .01 | .931 | ns |  | | -.19 | .21 |
|  |  |  | *Q-2* | .13 | .210 | ns |  |  | -.08 | .32 |
|  |  |  | *Q-3* | .24 | .008 | ** |  |  | .09 | .43 |
|  |  |  | *Q-4* | .15 | .127 | ns |  |  | -.05 | .34 |
|  | Crus I | b (paravermis) | **Avg.** | **.12** | **.226** | **ns** | **.226** | **ns** | **-.07** | **.31** |
|  |  |  | *Q-1* | -.11 | .266 | ns |  | | -.32 | .10 |
|  |  |  | *Q-2* | -.07 | .462 | ns |  |  | -.27 | .13 |
|  |  |  | *Q-3* | .20 | .042 | * |  |  | .02 | .38 |
|  |  |  | *Q-4* | .13 | .183 | ns |  |  | -.06 | .32 |
|  |  | c_1_ (cerebro-cerebellum) | **Avg.** | **.13** | **.210** | **ns** | **.226** | **ns** | **-.06** | **.31** |
|  |  |  | *Q-1* | -.17 | .084 | † |  | | -.36 | .02 |
|  |  |  | *Q-2* | -.02 | .862 | ns |  |  | -.21 | .19 |
|  |  |  | *Q-3* | .21 | .032 | * |  |  | .02 | .39 |
|  |  |  | *Q-4* | .12 | .221 | ns |  |  | -.07 | .31 |
|  | Crus II |  | **Avg.** | **.14** | **.178** | **ns** | **.226** | **ns** | **-.05** | **.32** |
|  |  |  | *Q-1* | .001 | .999 | ns |  | | -.22 | .23 |
|  |  |  | *Q-2* | -.09 | .402 | ns |  |  | -.28 | .12 |
|  |  |  | *Q-3* | .17 | .085 | † |  |  | -.01 | .35 |
|  |  |  | *Q-4* | .11 | .299 | ns |  |  | -.10 | .30 |
| R-R Mono-synaptic | Fastigial | | **Avg.** | **.25** | **.014** | ***** | **.036** | ***** | **.06** | **.41** |
|  |  |  | *Q-1* | .18 | .080 | † |  | | -.01 | .35 |
|  |  |  | *Q-2* | .25 | .011 | * |  |  | .06 | .43 |
|  |  |  | *Q-3* | .16 | .123 | ns |  |  | -.03 | .33 |
|  |  |  | *Q-4* | .11 | .260 | ns |  |  | -.08 | .30 |
|  | Interposed | | **Avg.** | **.23** | **.024** | ***** | **.036** | ***** | **.05** | **.39** |
|  |  |  | *Q-1* | .18 | .073 | † |  | | -.002 | .35 |
|  |  |  | *Q-2* | .23 | .022 | * |  |  | .04 | .41 |
|  |  |  | *Q-3* | .14 | .156 | ns |  |  | -.05 | .32 |
|  |  |  | *Q-4* | .08 | .416 | ns |  |  | -.12 | .28 |
|  | Dentate | | **Avg.** | **.17** | **.088** | **†** | **.088** | **†** | **-.02** | **.35** |
|  |  |  | *Q-1* | -.01 | .948 | ns |  | | -.21 | .19 |
|  |  |  | *Q-2* | .11 | .268 | ns |  |  | -.09 | .31 |
|  |  |  | *Q-3* | .21 | .040 | * |  |  | .02 | .38 |
|  |  |  | *Q-4* | .08 | .416 | ns |  |  | -.11 | .27 |

**Table S9.** Descriptive statistics for self-report measures under study.

| **Descriptive Statistics for Self-Report Measures** | | | | | | |
| --- | --- | --- | --- | --- | --- | --- |
| **Variable** | | | **Raw** | | **Percentile** | |
| **Assessment** | **Measure** | **Abbrev.** | **Mean** | **Std. Err.** | **Mean** | **Std. Err.** |
| **ASR** | *Anxiety/Depression* | Anx. Dep | 10.24 | 0.80 | 59.57 | 1.03 |
|  | *Internalizing* | Inter | 17.58 | 1.33 | 55.75 | 1.28 |
|  | *Social Withdrawal* | Soc. Withd | 3.43 | 0.30 | 56.49 | 0.79 |
|  | *Somaticizing* | Soma | 3.92 | 0.43 | 56.86 | 0.88 |
|  | *Attention Problems* | Attn | 8.24 | 0.56 | 57.70 | 0.81 |
|  | *Externalizing* | Exter | 11.20 | 0.81 | 52.07 | 0.92 |
|  | *Aggression* | Aggr | 4.94 | 0.42 | 54.15 | 0.60 |
| **NEO-FFI** | *Neuroticism* | N | 21.59 | 0.91 |  | |
|  | *Agreeableness* | A | 32.14 | 0.60 |  |  |
|  | *Conscientiousness* | C | 32.27 | 0.63 |  |  |
|  | *Extroversion* | E | 28.16 | 0.59 |  |  |

**Table S10.** Summary of descriptive statistics for each FA values of each tract and each quarter under investigation in the brain-behavior analyses. Note that these statistics are only for the right-seeded ipsilaterally traveling tract. N for all analyses is 101.

| **R-R Fractional Anisotropy Descriptive for Brain-Behavior Tracts** | | | | | |
| --- | --- | --- | --- | --- | --- |
| **Type** | **Lobule** | **Segment/DCN** | **Section** | **Mean** | **Std. Err.** |
| R-R Poly-synaptic | VI | a (vermis) | **Avg.** | **.44** | **.004** |
|  |  |  | *Q-1* | .37 | .003 |
|  |  |  | *Q-2* | .46 | .005 |
|  |  |  | *Q-3* | .50 | .006 |
|  |  |  | *Q-4* | .42 | .004 |
|  | Crus I | b (paravermis) | **Avg.** | **.45** | **.003** |
|  |  |  | *Q-1* | .36 | .003 |
|  |  |  | *Q-2* | .43 | .004 |
|  |  |  | *Q-3* | .55 | .006 |
|  |  |  | *Q-4* | .45 | .005 |
|  |  | c_1_ (cerebro-cerebellum) | **Avg.** | **.45** | **.003** |
|  |  |  | *Q-1* | .38 | .003 |
|  |  |  | *Q-2* | .41 | .004 |
|  |  |  | *Q-3* | .55 | .006 |
|  |  |  | *Q-4* | .46 | .005 |
|  | Crus II |  | **Avg.** | **.44** | **.003** |
|  |  |  | *Q-1* | .37 | .003 |
|  |  |  | *Q-2* | .38 | .003 |
|  |  |  | *Q-3* | .54 | .006 |
|  |  |  | *Q-4* | .47 | .005 |
| R-R Mono-synaptic | Fastigial | | **Avg.** | **.47** | **.005** |
|  |  |  | *Q-1* | .46 | .006 |
|  |  |  | *Q-2* | .53 | .007 |
|  |  |  | *Q-3* | .47 | .006 |
|  |  |  | *Q-4* | .41 | .004 |
|  | Interposed | | **Avg.** | **.50** | **.004** |
|  |  |  | *Q-1* | .50 | .005 |
|  |  |  | *Q-2* | .57 | .006 |
|  |  |  | *Q-3* | .51 | .007 |
|  |  |  | *Q-4* | .42 | .004 |
|  | Dentate | | **Avg.** | **.47** | **.004** |
|  |  |  | *Q-1* | .36 | .004 |
|  |  |  | *Q-2* | .53 | .006 |
|  |  |  | *Q-3* | .55 | .007 |
|  |  |  | *Q-4* | .45 | .005 |
| **L-L Fractional Anisotropy Descriptive for Brain-Behavior Tracts** | | | | | |
| **Type** | **Lobule** | **Segment/DCN** | **Section** | **Mean** | **Std. Err.** |
| L-L Poly-synaptic | VI | a (vermis) | **Avg.** | .40 | .004 |
|  |  |  | *Q-1* | .35 | .004 |
|  |  |  | *Q-2* | .41 | .005 |
|  |  |  | *Q-3* | .45 | .007 |
|  |  |  | *Q-4* | .39 | .004 |
|  | Crus I | b (paravermis) | **Avg.** | .43 | .003 |
|  |  |  | *Q-1* | .35 | .004 |
|  |  |  | *Q-2* | .43 | .003 |
|  |  |  | *Q-3* | .52 | .007 |
|  |  |  | *Q-4* | .43 | .005 |
|  |  | c_1_ (cerebro-cerebellum) | **Avg.** | .44 | .003 |
|  |  |  | *Q-1* | .36 | .003 |
|  |  |  | *Q-2* | .41 | .004 |
|  |  |  | *Q-3* | .53 | .007 |
|  |  |  | *Q-4* | .44 | .005 |
|  | Crus II |  | **Avg.** | .44 | .003 |
|  |  |  | *Q-1* | .37 | .003 |
|  |  |  | *Q-2* | .39 | .004 |
|  |  |  | *Q-3* | .53 | .006 |
|  |  |  | *Q-4* | .44 | .005 |
| L-L Mono-synaptic | Fastigial | | **Avg.** | .42 | .004 |
|  |  |  | *Q-1* | .38 | .005 |
|  |  |  | *Q-2* | .47 | .007 |
|  |  |  | *Q-3* | .44 | .006 |
|  |  |  | *Q-4* | .39 | .003 |
|  | Interposed | | **Avg.** | .47 | .005 |
|  |  |  | *Q-1* | .46 | .005 |
|  |  |  | *Q-2* | .53 | .007 |
|  |  |  | *Q-3* | .48 | .007 |
|  |  |  | *Q-4* | .40 | .003 |
|  | Dentate | | **Avg.** | .46 | .004 |
|  |  |  | *Q-1* | .36 | .004 |
|  |  |  | *Q-2* | .53 | .007 |
|  |  |  | *Q-3* | .52 | .008 |
|  |  |  | *Q-4* | .42 | .004 |

**Table S11.** Full correlation matrix with variables under study. Note that Spearman’s correlations are reported and there is an N of 101 for all analyses.

| **Variable** | | | **Stat** | Age | ASR | | | | | | | NEO-FFI | | | | Poly R-R FA | | | | Mono R-R FA | |
| --- | --- | --- | --- | --- | --- | --- | --- | --- | --- | --- | --- | --- | --- | --- | --- | --- | --- | --- | --- | --- | --- |
|  |  |  |  | 1. | 2. | 3. | 4. | 5. | 6. | 7. | 8. | 9. | 10. | 11. | 12. | 13. | 14. | 15. | 16. | 17. | 18. |
| Age | 1. | Age (Yrs) | *rho* |  | | | | | | | | | | | | | | | | | |
|  |  |  | *p* |  |  |  |  |  |  |  |  |  |  |  |  |  |  |  |  |  |  |
| ASR | 2. | Anx. Dep | *rho* | -.11 |  | | | | | | | | | | | | | | | | |
|  |  |  | *p* | .266 |  |  |  |  |  |  |  |  |  |  |  |  |  |  |  |  |  |
|  | 3. | Inter | *rho* | -.11 | .94 *** |  | | | | | | | | | | | | | | | |
|  |  |  | *p* | .290 | < .001 |  |  |  |  |  |  |  |  |  |  |  |  |  |  |  |  |
|  | 4. | Soc. With | *rho* | -.15 | .62 *** | .76 *** |  | | | | | | | | | | | | | | |
|  |  |  | *p* | .130 | < .001 | < .001 |  |  |  |  |  |  |  |  |  |  |  |  |  |  |  |
|  | 5. | Soma | *rho* | -.05 | .58 *** | .75 *** | .47 *** |  | | | | | | | | | | | | | |
|  |  |  | *p* | .628 | < .001 | < .001 | < .001 |  |  |  |  |  |  |  |  |  |  |  |  |  |  |
|  | 6. | Attn | *rho* | -.07 | .75 *** | .76 *** | .52 *** | .55 *** |  | | | | | | | | | | | | |
|  |  |  | *p* | .475 | < .001 | < .001 | < .001 | < .001 |  |  |  |  |  |  |  |  |  |  |  |  |  |
|  | 7. | Exter | *rho* | -.07 | .61 *** | .64 *** | .47 *** | .51 *** | .65 *** |  | | | | | | | | | | | |
|  |  |  | *p* | .518 | < .001 | < .001 | < .001 | < .001 | < .001 |  |  |  |  |  |  |  |  |  |  |  |  |
|  | 8. | Aggr | *rho* | -.08 | .69 *** | .67 *** | .51 *** | .45 *** | .61 *** | .82 *** |  | | | | | | | | | | |
|  |  |  | *p* | .402 | < .001 | < .001 | < .001 | < .001 | < .001 | < .001 |  |  |  |  |  |  |  |  |  |  |  |
| NEO-FFI | 9. | N | *rho* | -.10 | .88 *** | .81 *** | .53 *** | .49 *** | .62 *** | .53 *** | .65 *** |  | | | | | | | | | |
|  |  |  | *p* | .346 | < .001 | < .001 | < .001 | < .001 | < .001 | < .001 | < .001 |  |  |  |  |  |  |  |  |  |  |
|  | 10. | A | *rho* | .04 | -.03 | -.13 | -.35 *** | -.13 | -.07 | -.28 ** | -.23 * | -.11 |  | | | | | | | | |
|  |  |  | *p* | .673 | .738 | .191 | < .001 | .183 | .514 | .004 | .022 | .291 |  |  |  |  |  |  |  |  |  |
|  | 11. | C | *rho* | -.04 | -.40 *** | -.40 *** | 0.22 * | -.27 ** | -.70 *** | -.44 *** | -.34 *** | -.37 *** | .15 |  | | | | | | | |
|  |  |  | *p* | .667 | < .001 | < .001 | .028 | .006 | < .001 | < .001 | < .001 | < .001 | .144 |  |  |  |  |  |  |  |  |
|  | 12. | E | *rho* | -.07 | -.35 *** | -.41 *** | -.59 *** | -.17 † | -.30 ** | -.18 † | .24 * | -.33 *** | .29 ** | .24 * |  | | | | | | |
|  |  |  | *p* | .505 | < .001 | < .001 | < .001 | .087 | .002 | .066 | .014 | < .001 | .003 | .016 |  |  |  |  |  |  |  |
| Poly R-R FA | 13. | VI a | *rho* | -.01 | .28 ** | .28 ** | .15 | .28 ** | .20 * | .08 | .10 | .28 ** | .24 * | -.04 | -.06 |  | | | | | |
|  |  |  | *p* | .963 | .004 | .004 | .142 | .004 | .041 | .416 | .304 | .005 | .017 | .722 | .559 |  |  |  |  |  |  |
|  | 14. | Crus I b | *rho* | .09 | .23 * | .26 * | .19 † | .27 ** | .20 † | .02 | .02 | .27 ** | .17 | -.03 | -.15 | .79 *** |  | | | | |
|  |  |  | *p* | .362 | .020 | .010 | .064 | .007 | .050 | .841 | .811 | .006 | .100 | .807 | .129 | < .001 |  |  |  |  |  |
|  | 15. | Crus I c_1_ | *rho* | .17 | .18 † | .21 * | .20 * | .19 † | .20 † | .01 | -.002 | .22 * | .17 † | -.07 | -.11 | .67 *** | .85 *** |  | | | |
|  |  |  | *p* | .084† | .074 | .040 | .041 | .060 | .050 | .961 | .984 | .026 | .089 | .463 | .291 | < .001 | < .001 |  |  |  |  |
|  | 16. | Crus II c_2_ | *rho* | .08 | .22 * | .26 * | .23 * | .23 * | .23 * | .04 | .02 | .25 * | .17 † | -.14 | -.16 | .61 *** | .84 *** | .89 *** |  | | |
|  |  |  | *p* | .442 | .024 | .010 | .019 | .020 | .020 | .675 | .814 | .012 | .096 | .162 | .122 | < .001 | < .001 | < .001 |  |  |  |
| Mono R-R FA | 17. | FN | *rho* | .04 | .24 * | .23 * | .07 | .24 * | .15 | .05 | .06 | .25 * | .27 ** | .002 | -.03 | .94 *** | .76 *** | .65 *** | .61 *** |  | |
|  |  |  | *p* | .679 | .017 | .023 | .481 | .014 | .138 | .637 | .540 | .014 | .006 | .984 | .783 | < .001 | < .001 | < .001 | < .001 |  |  |
|  | 18. | IN | *rho* | .12 | .20 * | .20 * | .10 | .20 * | .18 † | -.01 | -.04 | .22 * | .26 * | -.06 | -.08 | .83 *** | .87 *** | .82 *** | .77 *** | .86 *** |  |
|  |  |  | *p* | .221 | .044 | .048 | .325 | .046 | .068 | .927 | .676 | .024 | .010 | .527 | .442 | < .001 | < .001 | < .001 | < .001 | < .001 |  |
|  | 19. | DN | *rho* | .13 | .14 | .17 † | .17 † | .15 | .13 | -.04 | -.02 | .18 † | .21 * | -.02 | -.09 | .62 *** | .82 *** | .89 *** | .89 *** | .64 *** | .85 *** |
|  |  |  | *p* | .184 | .159 | .098 | .090 | .139 | .181 | .696 | .871 | .074 | .033 | .882 | .392 | < .001 | < .001 | < .001 | < .001 | < .001 | < .001 |

† p < .10, * p < .05, ** p < .01, *** p < .001

**Table S12.** Heatmap of degree to which each quarter for the brain-behavior analyses is implicated. For polysynaptic tract investigations, the total possible number of times a single quarter may be implicated is four, while the total number of times a monosynaptic quarter may be implicated is three.

| **R-R Polysynaptic: 4 Tracts** | | | | | | | |
| --- | --- | --- | --- | --- | --- | --- | --- |
| **Assessment** | | **Variable** | **Abbrev.** | **Quarter** | | | |
|  |  |  |  | **Q1** | **Q2** | **Q3** | **Q4** |
| ASR | | *Anxiety/Depression* | Anx. Dep | 2 | 2 | 4 | 0 |
|  |  | *Internalizing* | Inter | 2 | 3 | 4 | 0 |
|  |  | *Social Withdrawal* | Soc. Withd | 4 | 2 | 1 | 0 |
|  |  | *Somaticizing* | Soma | 3 | 3 | 3 | 0 |
|  |  | *Attention Problems* | Attn | 1 | 2 | 2 | 0 |
| NEO-FFI | | *Neuroticism* | N | 0 | 2 | 4 | 0 |
|  |  | *Agreeableness* | A | 0 | 0 | 4 | 0 |
|  |  | *Neuroticism-Ctrl (Sex)* | N-Ctrl | 0 | 1 | 1 | 0 |
|  |  | *Agreeableness-Ctrl (Sex)* | A-Ctrl | 0 | 0 | 3 | 0 |
| **Total** | | | | **12** | **14** | **22** | **0** |
| **Total-Ctrl (Sex)** | | | | **12** | **13** | **18** | **0** |
| **R-R Monosynaptic: 3 Tracts** | | | | | | | |
| **Assessment** | **Variable** | | **Abbrev.** | **Quarter** | | | |
|  |  |  |  | **Q1** | **Q2** | **Q3** | **Q4** |
| ASR | *Anxiety/Depression* | | Anx. Dep | 1 | 2 | 1 | 0 |
|  | *Internalizing* | | Inter | 1 | 2 | 0 | 0 |
|  | *Social Withdrawal* | | Soc. Withd | 1 | 0 | 0 | 0 |
|  | *Somaticizing* | | Soma | 2 | 2 | 0 | 0 |
|  | *Attention Problems* | | Attn | 0 | 0 | 0 | 0 |
| NEO-FFI | *Neuroticism* | | N | 1 | 1 | 0 | 0 |
|  | *Agreeableness* | | A | 2 | 2 | 1 | 0 |
|  | *Neuroticism-Ctrl (Sex)* | | N-Ctrl | 0 | 1 | 0 | 0 |
|  | *Agreeableness-Ctrl (Sex)* | | A-Ctrl | 0 | 2 | 1 | 0 |
| **Total** | | | | **8** | **9** | **2** | **0** |
| **Total-Ctrl (Sex)** | | | | **5** | **9** | **2** | **0** |

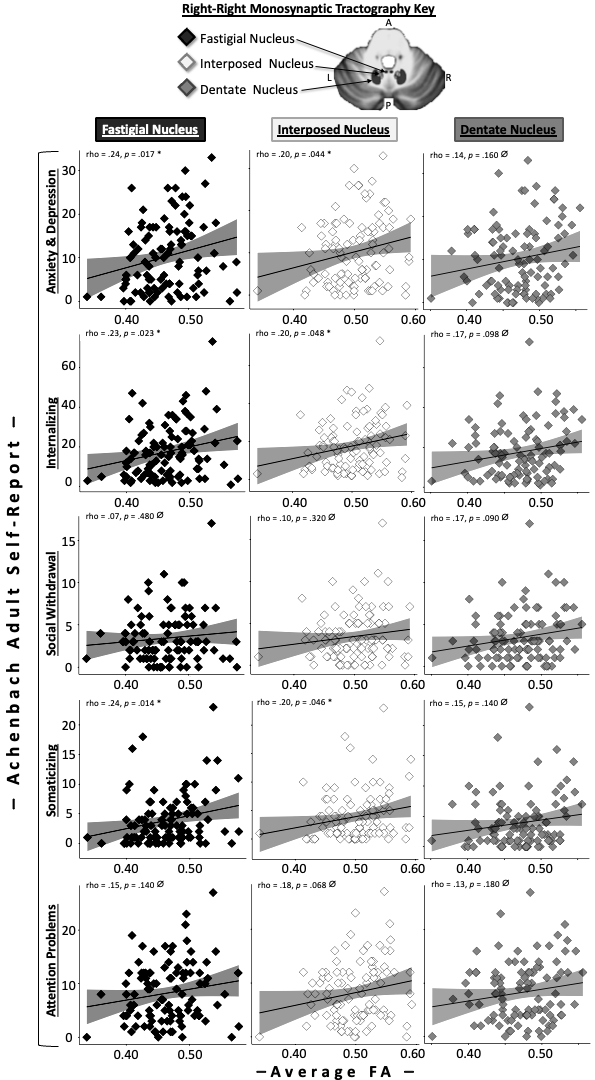

**Figure S2.** Summarizes results between monosynaptic tractography FA for each deep cerebellar nucleus traveling right-to-right and ASR measures. Note that FA is averaged across all 100 nodes for each tract, and the p-values are not corrected for multiple comparisons. **Significance flags:** ⌀ *p* > .05, * *p* < .05, ** *p* < .01

***Correlations with Achenbach Adult Self Report: Monosynaptic tracts***

*Anxiety and Depression.* For the monosynaptic tractographies, correlations only emerged for the tracts originating in the fastigial and interposed nuclei (findings became trends after FDR correction). For the quarter-wise analyses, Q1 and Q2 were implicated. Q3 and Q4 were never implicated. All findings held when controlling for multiple comparisons.

*Internalizing Symptoms*

Relations emerged for the fastigial and interposed monosynaptic tractography and internalizing symptoms, though they did not survive FDR correction. The only quarters implicated in these relations were Q1 and Q2.

*Social Withdrawal*

No tracts emerged as significant.

*Somatic Complaints*

For the monosynaptic tractographies, only the fastigial and interposed showed significant relations with somatic complaints; both Q1 and Q2 were implicated.

*Attention Problems*

No tracts emerged as significant.

*Neuroticism*

For the monosynaptic tractographies, only the fastigial and interposed tracts correlated with neuroticism, which survived FDR correction. Q1 and Q2 were implicated. Neither Q3 nor Q4 were ever implicated.

*Agreeableness*

For the monosynaptic tractographies, all tracts were significantly related to agreeableness. For the tracts originating in the fastigial and interposed, both Q1 and Q2 were implicated, whereas in the tract originating in the dentate, only Q3 was implicated.

***Table S13.*** Correlations with Achenbach Self-Report (ASR) subscales and Fractional Anisotropy (FA) values for the right-to-right travelling monosynaptic tracts with the highest densities.

| **Fractional Anisotropy Correlations: ASR Anxiety & Depression** | | | | | | | |
| --- | --- | --- | --- | --- | --- | --- | --- |
| **Deep Cerebellar Nucleus** | | **Section** | **Rho (*ρ*); *p*-value** | | ***p*-value FDR** | | **95% CI: Lower-Upper** |
| Fastigial | | **Avg.** | **.24**; p**=.017** | ***** | **.051 †** | | ***(.03*. *.42)*** |
|  |  | *Q-1* | .16; p**=**.107 | ns |  | |  |
|  |  | *Q-2* | .25; p**=**.011 | * |  |  |  |
|  |  | *Q-3* | .17; p**=**.096 | † |  |  |  |
|  |  | *Q-4* | .03; p**=**.778 | ns |  |  |  |
| Interposed | | **Avg.** | **.20**; p**=.044** | ***** | **.066 †** | | ***(.01*, *.38)*** |
|  |  | *Q-1* | .20; p**=**.044 | * |  | |  |
|  |  | *Q-2* | .22; p**=**.030 | * |  |  |  |
|  |  | *Q-3* | .13; p**=**.204 | ns |  |  |  |
|  |  | *Q-4* | .05; p**=**.606 | ns |  |  |  |
| Dentate | | **Avg.** | **.14**; p**=.159** | ns | **.159 †** | | **(-.07, .33)** |
|  |  | *Q-1* | .04; p**=**.707 | ns |  | |  |
|  |  | *Q-2* | .12; p**=**.231 | ns |  |  |  |
|  |  | *Q-3* | .14; p**=**.166 | ns |  |  |  |
|  |  | *Q-4* | .09; p**=**.348 | ns |  |  |  |
| **Fractional Anisotropy Correlations: ASR Internalizing Symptoms** | | | | | | | |
| **Deep Cerebellar Nucleus** | | **Section** | **Rho (*ρ*); *p*-value** | | ***p*-value FDR** | | **95% CI: Lower-Upper** |
| Fastigial | | **Avg.** | **.23**; p**=.023** | ***** | **.069 †** | | ***(.02*, *.42)*** |
|  |  | *Q-1* | .16; p**=**.110 | ns |  | |  |
|  |  | *Q-2* | .26; p**=**.010 | * |  |  |  |
|  |  | *Q-3* | .14; p**=**.155 | ns |  |  |  |
|  |  | *Q-4* | .01; p**=**.908 | ns |  |  |  |
| Interposed | | **Avg.** | **.20**; p**=.048** | ***** | **.072 †** | | ***(.001*, *.38)*** |
|  |  | *Q-1* | .20; p**=**.045 | * |  | |  |
|  |  | *Q-2* | .22; p**=**.026 | * |  |  |  |
|  |  | *Q-3* | .13; p**=**.200 | ns |  |  |  |
|  |  | *Q-4* | .04; p**=**.696 | ns |  |  |  |
| Dentate | | **Avg.** | **.17**; p**=.098** | **†** | **.098 †** | | **(-.04, .35)** |
|  |  | *Q-1* | .12; p**=**.240 | ns |  | |  |
|  |  | *Q-2* | .16; p**=**.117 | ns |  |  |  |
|  |  | *Q-3* | .15; p**=**.147 | ns |  |  |  |
|  |  | *Q-4* | .08; p**=**.459 | ns |  |  |  |
| **Fractional Anisotropy Correlations: ASR Social Withdrawal** | | | | | | | |
| **Deep Cerebellar Nucleus** | | **Section** | **Rho (*ρ*); *p*-value** | | ***p*-value FDR** | | **95% CI: Lower-Upper** |
| Fastigial | | **Avg.** | **.07**; p**=.481** | ns | **.481** ns | | **(-.12, .26)** |
|  |  | *Q-1* | .01; p**=**.900 | ns |  | |  |
|  |  | *Q-2* | .11; p**=**.256 | ns |  |  |  |
|  |  | *Q-3* | .03; p**=**.734 | ns |  |  |  |
|  |  | *Q-4* | -.05; p**=**.605 | ns |  |  |  |
| Interposed | | **Avg.** | **.10**; p**=.325** | ns | **.481** ns | | **(-.09, .29)** |
|  |  | *Q-1* | .08; p**=**.439 | ns |  | |  |
|  |  | *Q-2* | .11; p**=**.261 | ns |  |  |  |
|  |  | *Q-3* | .08; p**=**.436 | ns |  |  |  |
|  |  | *Q-4* | .001;p**=**.996 | ns |  |  |  |
| Dentate | | **Avg.** | **.17**; p**=.090** | **†** | **.270** ns | | **(-.04, .36)** |
|  |  | *Q-1* | .24; p**=**.018 | * |  | |  |
|  |  | *Q-2* | .15; p**=**.125 | ns |  |  |  |
|  |  | *Q-3* | .11; p**=**.268 | ns |  |  |  |
|  |  | *Q-4* | .05; p=.597 | ns |  |  |  |
| **Fractional Anisotropy Correlations: ASR Somaticizing Symptoms** | | | | | | | |
| **Deep Cerebellar Nucleus** | | **Section** | **Rho (*ρ*); *p*-value** | | ***p*-value FDR** | **95% CI: Lower-Upper** | |
| Fastigial | | **Avg.** | **.24**; p**=.014** | ***** | **.042 *** | **(.03, .45)** | |
|  |  | *Q-1* | .26; p**=**.008 | ** |  |  | |
|  |  | *Q-2* | .26; p**=**.008 | ** |  |  |  |
|  |  | *Q-3* | .10; p**=**.344 | ns |  |  |  |
|  |  | *Q-4* | -.01; p**=**.918 | ns |  |  |  |
| Interposed | | **Avg.** | **.20**; p**=.046** | ***** | **.069 †** | **(-.01, .39)** | |
|  |  | *Q-1* | .23; p**=**.019 | * |  |  | |
|  |  | *Q-2* | .24; p**=**.017 | * |  |  |  |
|  |  | *Q-3* | .11; p**=**.290 | ns |  |  |  |
|  |  | *Q-4* | -.01; p**=**.945 | ns |  |  |  |
| Dentate | | **Avg.** | **.15**; p**=.139** | ns | **.139** ns | **(-.06, .35)** | |
|  |  | *Q-1* | .15; p**=**.143 | ns |  |  | |
|  |  | *Q-2* | .15; p**=**.125 | ns |  |  |  |
|  |  | *Q-3* | .14; p**=**.164 | ns |  |  |  |
|  |  | *Q-4* | -.01; p**=**.940 | ns |  |  |  |
| **Fractional Anisotropy Correlations: ASR Attention Problems** | | | | | | | |
| **Deep Cerebellar Nucleus** | **Section** | | **Rho (*ρ*); *p*-value** | | ***p*-value FDR** | **95% CI: Lower-Upper** | |
| Fastigial | **Avg.** | | **.15**; p**=.138** | ns | **.181** ns | **(-.0, -.33)** | |
|  | *Q-1* | | .10; p**=**.310 | ns |  |  | |
|  | *Q-2* | | .17; p**=**.100 | ns |  |  |  |
|  | *Q-3* | | .09; p**=**.376 | ns |  |  |  |
|  | *Q-4* | | .004; p**=**.968 | ns |  |  |  |
| Interposed | **Avg.** | | **.18**; p**=.068** | **†** | **.181** ns | **(-.01, .36)** | |
|  | *Q-1* | | .18; p**=**.074 | † |  |  | |
|  | *Q-2* | | .19; p**=**.054 | † |  |  |  |
|  | *Q-3* | | .11; p**=**.278 | ns |  |  |  |
|  | *Q-4* | | .01; p**=**.946 | ns |  |  |  |
| Dentate | **Avg.** | | **.13**; p**=.181** | ns | **.181** ns | **(-.07, .33)** | |
|  | *Q-1* | | .12; p**=**.246 | ns |  |  | |
|  | *Q-2* | | .09; p**=**.358 | ns |  |  |  |
|  | *Q-3* | | .15; p**=**.133 | ns |  |  |  |
|  | *Q-4* | | .05; p**=**.652 | ns |  |  |  |

**Significance Flags**: † p < .10, * p < .05, ** p < .01, *** p < .001
